# ZCWPW1 organizes telomeric architecture to drive meiotic chromosome movements

**DOI:** 10.64898/2026.01.30.702899

**Authors:** Wenxin Xie, Manjunath Gowder, Dominic Bazzano, Adrienne Niederriter Shami, Aastha Pandey, Melissa Frasca, Lakshmi Paniker, Luciana Previato de Almeida, Binod Kumar Bharati, Jing Liang, Attila Tóth, Morgan DeSantis, Jayakrishnan Nandakumar, Evgeny Nudler, Shyamal Mosalaganti, Roberto Pezza, Francesca Cole, Saher Sue Hammoud

## Abstract

Meiotic homolog pairing relies on programmed DNA recombination and large-scale chromosome movements, yet, how these genetic and mechanical events are coordinated remains unclear. ZCWPW1 is a histone reader that recognizes PRDM9-deposited chromatin marks. We identify an unexpected role for ZCWPW1 as a regulator of rapid prophase movements (RPMs). Using super-resolution imaging, we show that ZCWPW1 is strongly enriched at subtelomeric regions of mouse spermatocytes, where it stabilizes TRF1, LINC complex components, dynein, and meiosis-specific cohesin (STAG3). Loss of ZCWPW1 disrupts telomere architecture, weakens telomere–LINC– motor coupling, and abolishes chromosome movement, leading to defective synapsis and pairing, and persistence of DSBs. These defects are more severe than, and mechanistically independent of, those observed in *Prdm9*^−*/*−^ spermatocytes. Together, our findings reveal that ZCWPW1 acts independently of PRDM9 as a chromatin-based intranuclear regulator of telomere architecture and telomere-led chromosome movements, thereby linking telomeric chromatin state to nuclear force transmission required for faithful meiotic progression.

**Significance Statement:** Meiotic pairing requires recombination and telomere-led chromosome movements, yet no chromatin factor has been shown to regulate both. We identify ZCWPW1 as the first chromatin-based regulator of rapid prophase movements. ZCWPW1 organizes telomeric chromatin and promotes cohesin and motor assembly at telomeres that for force transmission across the nuclear envelope. Loss of ZCWPW1 disrupts the telomere-nuclear envelope mechanical coupling, impairing motion and altering recombination. This function doesn’t rely on PRDM9 despite their co-evolution and co-expression, challenging the prevailing view that ZCWPW1 only acts downstream of PRDM9 in DNA repair. Our findings show that chromatin readers can function as structural regulators of genome organization, revealing a conserved mechanism integrating chromosome structure, motion, and repair to ensure proper pairing and fertility.

## Text Introduction

In sexually reproducing organisms, homologous chromosome pairing is an essential and evolutionarily conserved feature of meiosis. In many organisms including mammals, pairing is tightly associated with recombination-dependent double-strand break (DSB) repair between homologs and ensures faithful chromosome segregation essential for healthy haploid gametes. Despite its fundamental importance, the mechanisms governing the pairing process are poorly understood. However, two core and conserved regulators consistently emerge: meiotic recombination and large-scale chromosome movements.

In mammals, recombination sites are specified by the histone methyltransferase PRDM9, which deposits H3K4me3 and H3K36me3 marks at recombination hotspots on both homologs (1– 3). These modifications help recruit the DSB machinery. Efficient repair of DSBs by homologous recombination requires the timely association of homologous chromosomes within the nucleus, while recombination intermediates in turn promote chromosome-specific pairing and synapsis.

Proper homologous chromosome interactions are closely associated with dynamic telomere-led chromosomal movements, referred to as rapid prophase movements (RPMs) (4–6). RPMs are generated in the cytoplasm by molecular motors walking on microtubules associated with the nuclear membrane. These forces are transmitted to chromosome ends attached to the nuclear envelope (NE) through the NE-spanning Linker of Nucleoskeleton and Cytoskeleton (LINC) complex composed of SUN/KASH (4, 7–12). In most species, telomeres serve as both anchor points and mechanical handles for pulling forces exerted by cytoskeletal motors. In mammals, the TERB1–TERB2–MAJIN (TTM) complex tethers telomeres to the NE (7, 13, 14), which in turn interacts with the LINC complex and indirectly with cytoskeletal motors to allow movement. Although RPMs and recombination rely on distinct molecular machinery, these processes are tightly interconnected across many species (12, 15, 16). Perturbations in RPMs impair DNA repair efficiency (12, 15, 16), while recombination and synapsis mutants can also impair RPMs or bouquet exit (17, 18). This coupling of chromosome movement and repair is not unique to meiosis but is also observed in mitotic cells in many organisms (19–23). Despite this functional interdependence, the molecular players that directly connect chromatin-based recombination events with mechanisms of chromosome movements remain largely unknown.

Here, we identify the meiosis-specific chromatin reader ZCWPW1 as a key regulator of recombination and chromosome dynamics in mammalian meiosis. ZCWPW1 contains two epigenetic reader domains: the zf-CW domain that binds H3K4me3, and the PWWP domain that recognizes H3K36me3 (24), both of which are highly conserved in vertebrates (25, 26). While each domain is individually found in diverse chromatin regulators such as histone modifiers, DNA methylases, and repair proteins, their combination within ZCWPW1 makes it uniquely poised to recognize the dual histone H3K4/K36 marks deposited by PRDM9 at recombination hotspots (25– 27). Consistent with this, ZCWPW1 has co-evolved with PRDM9 (25, 26, 28), is strongly recruited to PRDM9 sites genome-wide (25–27), and is essential for male meiosis. Unexpectedly, we also find that ZCWPW1 localizes to subtelomeric regions at the nuclear periphery of mouse spermatocytes, a pattern that is maintained even in the absence of PRDM9, suggesting an additional PRDM9-independent role. Loss of ZCWPW1 causes severe meiotic defects, including delayed homolog pairing, loss of RPMs, and weakened LINC complex and dynein recruitment - phenotypes more severe than in *Prdm9*^−*/*−^ mice. Moreover, *Zcwpw1*^−*/*−^ spermatocytes have abnormally decompacted telomeres and reduced levels of cohesin. These changes weaken the mechanical coupling between telomeres and the LINC complex and diminish the efficiency of dynein-driven chromosome movement. As a result, we find that synapsis and homolog pairing are significantly impaired in *Zcwpw1*^−*/*−^ spermatocytes. Together, these findings indicate that although ZCWPW1 likely co-evolved with PRDM9 to support recombination, it has acquired a PRDM9-independent role that also promotes homolog pairing as a chromatin-based intranuclear regulator of processive chromosome movements.

## Results

### ZCWPW1 loss causes more severe defects in meiotic DSB repair and synapsis than PRDM9 loss

To examine the role of ZCWPW1 in mammalian meiosis, we used CRISPR-Cas9 to generate *Zcwpw1*-knockout (KO) mice (***SI Appendix*, Fig. S1A**,**B**) in two genetic backgrounds: C57BL/6J (B6) and DBA/2J (DBA) strains. We confirmed loss of the ZCWPW1 protein in *Zcwpw1*^−*/*−^ testes by western blot (***SI Appendix*, Fig. S1C**) and immunofluorescence (***SI Appendix*, Fig. S1D**) and noted that wildtype (*Zcwpw1*^*+/+*^ or WT) and heterozygous (*Zcwpw1*^*+/*−^ or HET) mice were phenotypically indistinguishable, despite modest differences in ZCWPW1 protein levels (***SI Appendix*, Fig. S1C**). Therefore, in some cases we pooled WT and HET samples for analysis and these results are referred as WT* (***SI Appendix*, Fig. S1C–F**).

As previously reported, the testis-to-body weight ratios in *Zcwpw1*^−*/*−^ males were indistinguishable from WT during juvenile stages but were significantly reduced in adulthood (***SI Appendix*, Fig. S1E–G**). Histological analysis of *Zcwpw1*^−*/*−^ *males* revealed increased germ cell apoptosis (***SI Appendix*, Fig. S1J**) and complete absence of post-meiotic germ cells in *testes* (***SI Appendix*, Fig. S1H**,**I**), and depletion of sperm in the epididymis (***SI Appendix*, Fig. S1H**,**I**), a phenotype that closely resembles *Prdm9*^−*/*−^ males (***SI Appendix*, Fig. S1K**). The *Zcwpw1*^−*/*−^ spermatocytes had meiotic arrest before the mid-pachynema stage based on the extent of synapsis and H1t expression (hereafter referred as pachynema-like, ***SI Appendix*, Fig. S2A**,**B**). In contrast, *Zcwpw1*^−*/*−^ *females* remained fertile (data not shown). Together, these findings confirm that ZCWPW1 is required for male meiosis in two genetic backgrounds. Unless specified, we used B6 male mice for subsequent experiments.

A key hallmark of meiotic prophase I is the programmed induction of DSBs by the topoisomerase-like enzyme SPO11 primarily at hotspots. The number and genomic distribution of DSBs were unaffected in the *Zcwpw1*^−*/*−^, as revealed by END-seq and DMC1 ChIP-seq (25, 26), and here when comparing the number of RAD51 and RPA2 foci in leptonema across genotypes (***SI Appendix*, Fig. S2K**,**L**). However, these breaks failed to repair in the absence of ZCWPW1. Specifically, markers of unrepaired DSBs including γH2AX (***SI Appendix*, Fig. S2C**), RPA2, and RAD51 (***SI Appendix*, Fig. S2I–L**), peaked in early zygonema and persisted abnormally into pachynema-like cells. Notably, the number of RAD51 foci was significantly higher in later stages from *Zcwpw1*^−*/*−^ mice compared to *Prdm9*^−*/*−^ mice (mean ± s.d.: WT, 30.05 ± 18.68; *Zcwpw1*^−*/*−^, 109.6 ± 34.27; *Prdm9*^−*/*−^, 61.82 ± 25.60; ***SI Appendix*, Fig. S2K**), suggesting a more profound defect in DSB repair in *Zcwpw1*^−*/*−^. While the RPA2 foci did not follow the same quantitative trends as RAD51 across stages, the later RPA2 patterns likely reflect its broader roles in recombination dynamics, e.g. early resection intermediates and D-loops (30).

Given the persistence of DSBs, we next examined the synaptonemal complex (SC) assembly, which is known to stabilize homolog interactions, promote proper maturation of crossover intermediates, and suppress continued DSB accumulation during meiotic prophase. Consistent with previous reports (25–27, 29), *Zcwpw1*^−*/*−^ spermatocytes displayed severe defects in synapsis. On average, less than half of the chromosomes completed synapsis, as indicated by complete overlap of SYCP1 and SYCP3, which label the transverse filaments and lateral elements of the SC, respectively (mean ± s.d.: 8.306 ± 2.646, ***SI Appendix*, Fig. S2D**,**E**). As expected, the unsynapsed chromosomal axes retained HORMAD1 (***SI Appendix*, Fig. S2D**,**F**). To further assess SC formation and compare synapsis efficiency across genotypes, we performed super-resolution imaging of SYCP1 and SYCP3 in WT, *Zcwpw1*^−*/*−^, and *Prdm9*^−*/*−^ spermatocytes. This analysis revealed that both *Zcwpw1*^−*/*−^ and *Prdm9*^−*/*−^ spermatocytes displayed a high frequency of asynapsis and nonhomologous (heterologous) synapsis, but these abnormalities were more pronounced in *Zcwpw1*^−*/*−^ (***SI Appendix*, Fig. S2G**,**H**). Together, these findings demonstrate that both synapsis and DSB repair are more severely compromised in the absence of ZCWPW1 than PRDM9.

### ZCWPW1 is recruited to meiotic DSBs and promotes recombination resolution

Given that ZCWPW1 is a histone methylation reader proposed to act downstream of PRDM9 at recombination hotspots, we asked whether ZCWPW1 enriches at DSB sites *in vivo* at the single cell level. To examine this, we co-stained testis cross-sections from WT and *Prdm9*^−*/*−^ for ZCWPW1 and RAD51. In WT spermatocytes, ZCWPW1 localized along chromosome axes and overlapped with RAD51 foci in 73.7% of cases (**Fig. 1A,B**). This colocalization of ZCWPW1 and RAD51 was largely maintained in *Prdm9*^−*/*−^ (68.3%; **Fig. 1B**); indicating that ZCWPW1 is recruited to meiotic DSBs independently of PRDM9. These findings are consistent with ChIP-seq and CUT&RUN data showing co-occupancy of ZCWPW1 and DMC1 at recombination hotspots in WT spermatocytes (25, 27), and together support a direct role for ZCWPW1 at meiotic DSB repair.

**Figure 1.**
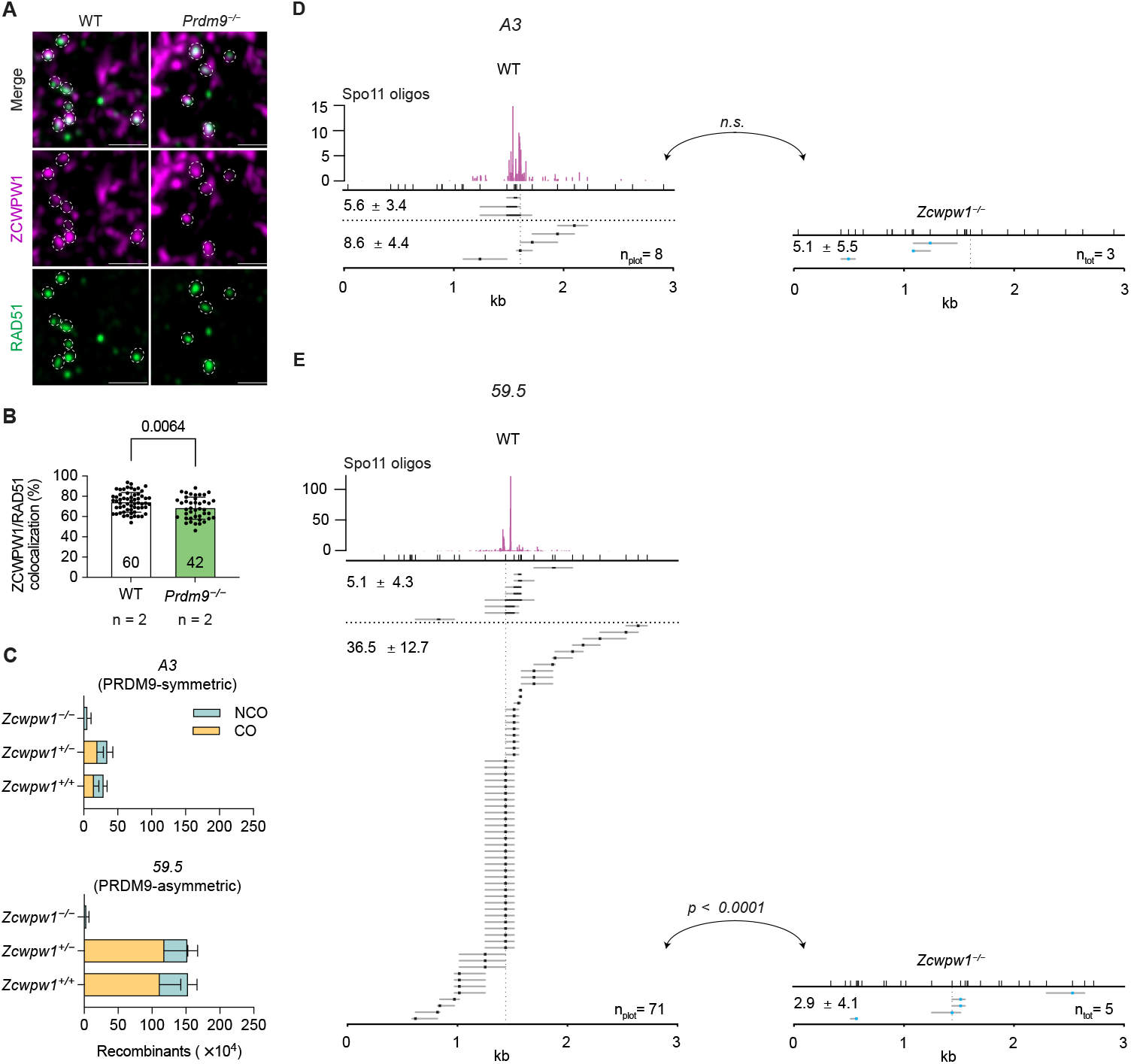
ZCWPW1 localizes to recombination sites and *Zcwpw1*^−*/*−^ spermatocytes are deficient in recombination. (*A*) Immunofluorescence of ZCWPW1 and RAD51 in WT and *Prdm9*^−*/*−^ spermatocytes. Scale bars: 1 μm. (*B*) Quantification of RAD51 foci colocalizing with ZCWPW1 (mean ± s.d.; Welch’s t-test). The number of cells analyzed per group are indicated below each bar. (*C*) Quantification of COs and NCOs at *A3* and *59*.*5* hotpsots in *Zcwpw1*^*+/+*^ (*A3*: n = 6; *59*.*5*: n = 11), *Zcwpw1*^*+/*−^ (n = 3 for both *A3* and *59*.*5*), and *Zcwpw1*^−*/*−^ (*A3*: n = 3; *59*.*5*: n = 5) male mice. (*D–E*) Representative gene co-conversions and singleton noncrossovers for the *A3* (*D*) and *59*.*5* (*E*) hotspots in *Zcwpw1*^*+/+*^ and *Zcwpw1*^−*/*−^ 4C spermatocytes. Top track: SPO11 oligonucleotide map in reads per million in magenta (64). The colored square plots the minimum, and the grey line plots the maximum possible gene conversion tract. The top x-axis marks the polymorphisms analyzed, and the bottom x-axis marks the relative length and position within the hotspot, starting at 0 kb. The vertical dotted line marks the center of the hotspot, and the horizontal dashed line separates the singleton noncrossover and co-conversion events. The total number of events plotted (n_plot_) or the total number of observed (n_tot_) is shown at the bottom right. The total frequency ± s.d. of singleton noncrossover and co-conversion events per 10,000 molecules is shown at the top left. *P* values were calculated using Chi-square with Yates correction, 2-tailed.

To determine the role of ZCWPW1 in DSB repair and recombination resolution, we monitored both crossover and noncrossover recombination in 4C spermatocytes isolated from *Zcwpw1*^−*/*−^ and WT B6 × DBA F1 hybrid mice (***SI Appendix*, Fig. S3A**,**B**). We analyzed two well-characterized PRDM9-directed hotspots: (1) *A3* on chr.1 (a PRDM9-symmetric site with both parental chromosomes having an equal likelihood of receiving DSBs (31)), and (2) *59*.*5* on chr.19 (a PRDM9-asymmetric site with over 90% of DSBs incurred on the B6 chromosome (32)). The total recombination events were significantly decreased in *Zcwpw1*^−*/*−^ 4C spermatocytes as compared to *Zcwpw1*^*+/+*^ and *Zcwpw1*^*+/*−^. Importantly, these defects in recombination were independent of PRDM9 symmetry (**Fig. 1C**). Furthermore, *Zcwpw1*^−*/*−^ had a complete loss of crossovers at both *59*.*5* and *A3* hotspots (**Fig. 1C**), consistent with the absence of MLH1 foci previously described (26, 27, 29). Curiously, while singleton noncrossovers that involve gene conversion of only a single polymorphism were still detected at lower frequencies in *Zcwpw1*^−*/*−^ 4C spermatocytes, we did not find any co-converted noncrossovers that involve gene conversion of multiple and often adjacent polymorphisms at either hotspot (**Fig. 1D,E**). Therefore, although factors responsible for crossover designation (i.e. MSH4 and RNF212) were previously shown to localize to recombination sites (27), interhomolog alignment may be unstable and only support repair via noncrossover pathways such as Synthesis-Dependent Strand Annealing (SDSA) (which mostly produce singleton noncrossovers). Alternatively, reduced chromatin accessibility at meiotic hotspots due to loss of H3K9ac in *Zcwpw1*^−*/*−^ spermatocytes (33) can bias DSB repair away from stable crossover formation and instead favor SDSA. Taken together, bulk sequencing and single cell observations suggest a direct role for ZCWPW1 in DSB repair and recombination events.

### ZCWPW1 is required for efficient and timely homolog pairing and synapsis

To investigate the basis of aberrant synapsis and inefficient recombination in *Zcwpw1*^−*/*−^, we combined whole chromosome DNA-PAINT with immunofluorescence to evaluate the pairing frequency of chr.1 and chr.19 homologs. These chromosomes differ in length, chr.1 is the largest and chr.19 the smallest autosome in mice, and each harbors a recombination hotspot used in our earlier assays (**Fig. 2A**). This setup allowed us to assess how ZCWPW1 affects pairing across chromosomes of different lengths. To quantify pairing, we measured the percentage of spermatocyte nuclei in which chr.1 or chr.19 homologs appeared as a single diffraction-limited domain (**Fig. 2B**).

**Figure 2.**
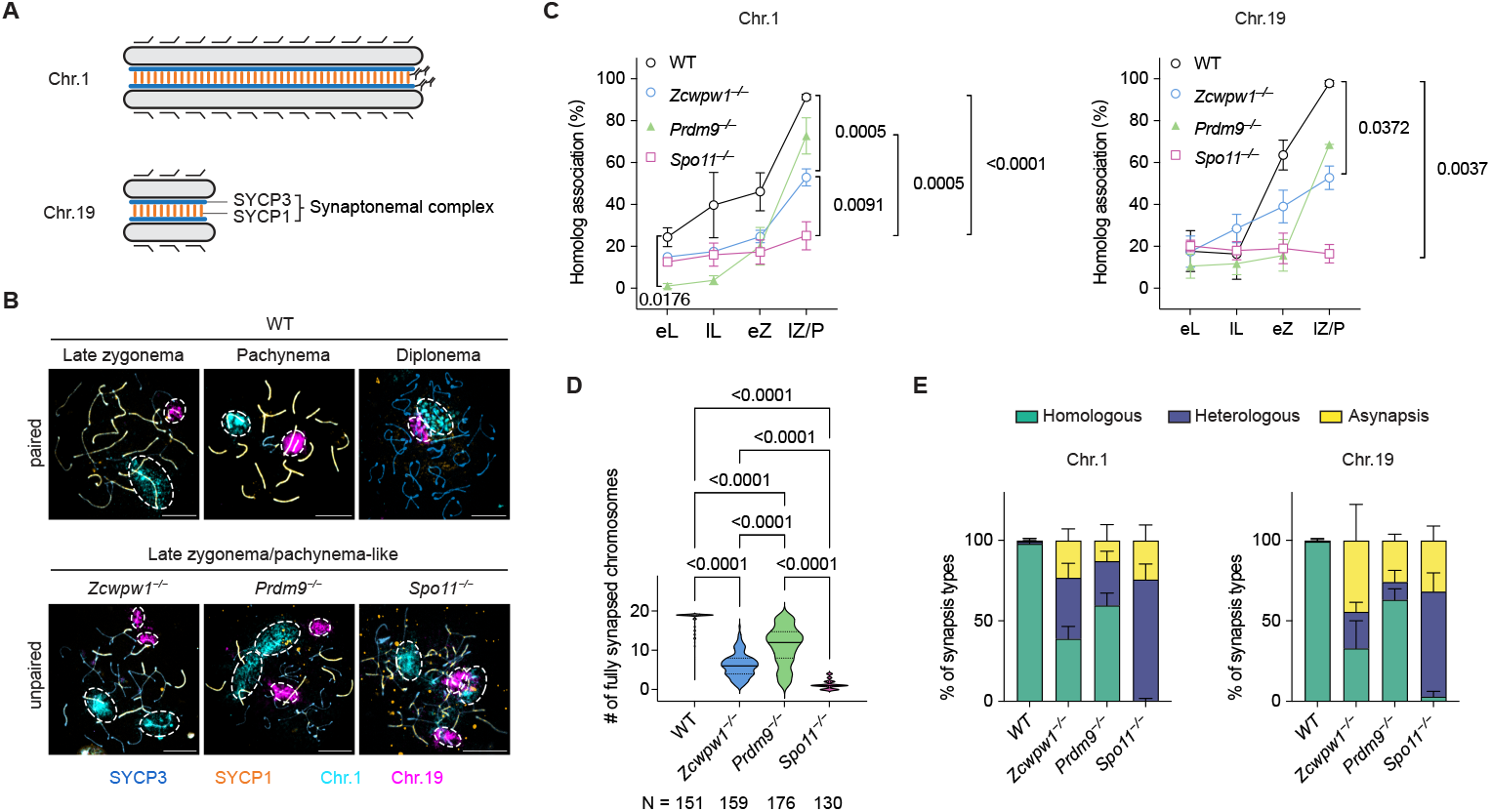
ZCWPW1 is required for the timely and efficient homolog pairing. (*A*) Schematic of chromosome painting for chr.1 and chr.19 combined with synaptonemal complex labeling to assess pairing and synapsis. (*B*) Representative images of pairing and synapsis in meiotic prophase I. Dashed circle, a single cloud of chr.1 or chr.19. Scale bars,10 μm. (*C*) Dynamics of pairing of chr.1 and chr.19 homologs during meiotic prophase I of WT (n = 5), *Zcwpw1*^−*/*−^ (n = 7), *Prdm9*^−*/*−^ (n = 3), and *Spo11*^−*/*−^ (n = 3) mice. eL: early leptonema; lL: late leptonema; eZ: early zygonema; lZ/P: late zygonema, pachynema or pachynema-like. Data are mean ± s.e.m. n: biological replicates; N: cells analyzed per group. *P* = 0.0176 (WT and *Prdm9*^−*/*−^ at eL), and *p* values at lZ/P from Kruskal–Wallis tests. (*D*) Quantification of the number of fully synapsed autosomes in zygonema and pachynema/pachynema-like, n = 3 for each group. Data are mean ± s.d. *P* values from Kruskal– Wallis test. (*E*) Quantification of different synapsis types for chr.1 and chr.19 in late zygonema and pachynema or pachynema-like of WT, *Zcwpw1*^−*/*−^, *Prdm9*^−*/*−^, and *Spo11*^−*/*−^ mice, n = 3 for each group.

In wild-type (WT) spermatocytes, pairing of chr.1 and chr.19 was sometimes detected in early leptonema (∼20%) and steadily increased across prophase I, reaching 89.6% and 97.4%, respectively, in late zygonema and pachynema (**Fig. 2B,C**). In contrast, *Zcwpw1*^−*/*−^ spermatocytes had a similar early pairing profile but progressed more slowly and plateaued at ∼53% for both chromosomes by pachynema. To contextualize the phenotype of ZCWPW1 loss with other meiotic mutants, we examined *Prdm9*^−*/*−^ and *Spo11*^−*/*−^ mutants. *Prdm9*^−*/*−^ spermatocytes, initiated pairing at lower frequencies than WT in leptonema but quickly increased to ∼73% for chr.1 and ∼74% for chr.19 by pachynema - intermediate levels between WT and *Zcwpw1*^−*/*−^ (**Fig. 2C**). In contrast, *Spo11*^−*/*−^ spermatocytes, which lack programmed meiotic DSBs, had only basal pairing throughout prophase I (34) (chr.1: 24.4%; chr.19: 16.5%). This result is in agreement with the essential role of recombination intermediates in stabilizing homologous chromosome interactions (35, 36).

Consistent with these pairing frequencies, the number of fully synapsed chromosomes - visualized via immunofluorescence for synaptonemal complex components - was significantly reduced in *Zcwpw1*^−*/*−^ relative to WT and *Prdm9*^−*/*−^, but greater than in *Spo11*^−*/*−^ (**Fig. 2D**). Combining DNA-PAINT with synaptonemal complex staining further revealed increased asynapsis and nonhomologous synapsis in *Zcwpw1*^−*/*−^ compared to *Prdm9*^−*/*−^ spermatocytes (**Fig. 2E**).

Taken together, our data demonstrate that ZCWPW1 is required for the timely and efficient progression of homolog pairing and synapsis, independent of chromosome length. Notably, the loss of ZCWPW1 leads to a more severe defect than *Prdm9* deficiency: fewer homologous chromosomes pair in *Zcwpw1*^−*/*−^ than in *Prdm9*^−*/*−^ spermatocytes. Therefore, while both factors are essential, ZCWPW1 likely promotes pairing in a PRDM9-independent manner.

### ZCWPW1 localizes to the inner nuclear envelope and near telomeres

To examine why ZCWPW1 causes a more severe meiotic defects than *Prdm9*^−*/*−^, we examined the expression and localization of both proteins in mouse spermatocytes. Cross-species scRNA-seq from human, mouse, and macaque testes revealed a conserved onset of *Zcwpw1* transcription between type B spermatogonia and early preleptotene spermatocytes (37). At the protein level, ZCWPW1 was detected in human testes (***SI Appendix*, Fig. S4A**) and in mouse type B spermatogonia and primary spermatocytes: spanning pre-leptonema to mid-pachynema, with peak abundance in zygonema and early pachynema (**Fig. 3A**). This temporal expression pattern mirrors staining patterns observed in spermatocyte spreads (26) and by immunoblots of juvenile testes (***SI Appendix*, Fig. S4B**). In addition to the diffuse nuclear staining, which potentially reflects the H3K4me3/H3K36me3 reading function of ZCWPW1, we observed a thickened nuclear rim staining from early zygonema to early pachynema (**Fig. 3A**). This localization pattern is unique to ZCWPW1 and was not observed for PRDM9 (***SI Appendix*, Fig. S4C**). Using super-resolution microscopy, we confirmed that ZCWPW1 resides beneath the nuclear lamina (***SI Appendix*, Fig. S4D**) and nuclear pores (Nup153) (***SI Appendix*, Fig. S4E**), a configuration that persists in *Prdm9*^−*/*−^ spermatocytes. Curiously, this peripheral ZCWPW1 enrichment is excluded from but adjacent to the DAPI-dense heterochromatin (***SI Appendix*, Fig. S4E**) and specifically enriches in telomere-adjacent (subtelomeric) regions during zygonema and pachynema (**Fig. 3B,C**). Together, these observations reveal a unique ZCWPW1-specific enrichment at telomere-adjacent regions, suggesting a role in regulating meiotic telomere architecture and function.

**Figure 3.**
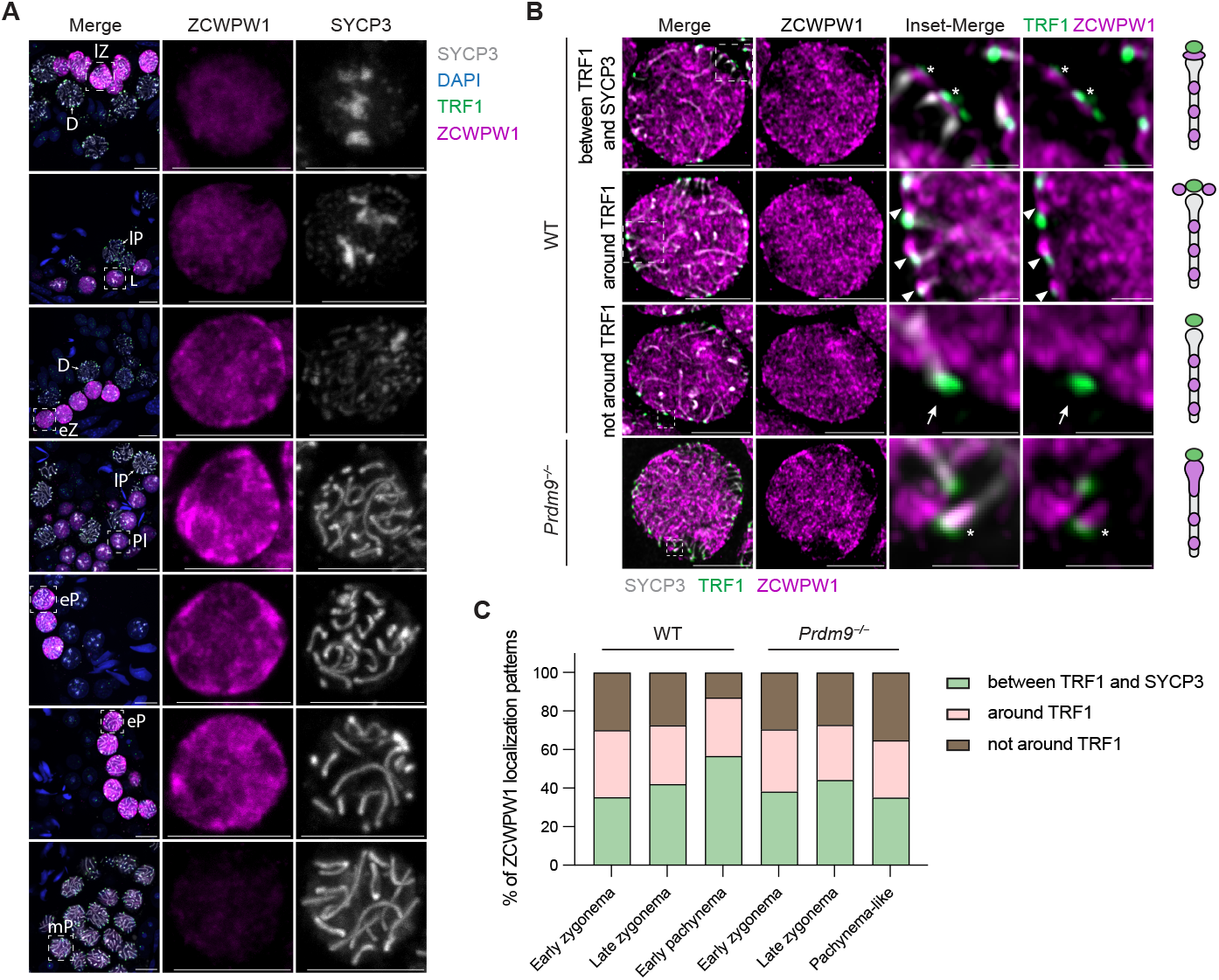
ZCWPW1 is enriched at the nuclear periphery and near telomeres. (*A*) Immunofluorescence of SYCP3, and ZCWPW1, and TRF1 on testis cross-sections from adult WT mice showing ZCWPW1 localization during meiotic prophase I. Nuclei were counterstained by 4′,6-diamidino-2-phenylindole (DAPI). Insets show separate channels of cells highlighted with dashed squares. Meiotic stages: Pl, preleptonema; L, leptonema; eZ, early zygonema; lZ, late zygonema; eP, early pachynema; mP, mid pachynema; lP, late pachynema; D, diplonema. Scale bars, 10 μm. Airyscan images of SYCP3, ZCWPW1, and TRF1 in zygotene spermatocytes from adult WT and *Prdm9*^−*/*−^ mice. Insets (dashed squares) highlight distinct ZCWPW1 localization patterns: (top) ZCWPW1 between TRF1 and SYCP3 ends (asterisks); (middle) ZCWPW1 adjacent to TRF1 at the nuclear periphery (arrowheads); (bottom) ZCWPW1 positioned apart from TRF1 (arrows). Scale bars, 5 μm (main images); 1 μm (insets). (*C*) Quantification of ZCWPW1 localization patterns in early zygonema, late zygonema, and early pachynema of WT and *Prdm9*^−*/*−^ mice.

### ZCWPW1 has a PRDM9-independent role in rapid prophase movement

The enrichment of ZCWPW1 at the nuclear periphery, coupled with its association to telomeres, prompted us to test whether it regulates telomere-led rapid prophase movements (RPMs). RPMs include both whole-nucleus rotations and individual chromosomal movements that are essential for efficient homology search and pairing in early meiotic prophase I (4–7, 18). RPMs require the localization of telomeres to the nuclear envelope, where they are connected to cytoplasmic dynein motors via the LINC complex. This connection allows mechanical forces generated by motor proteins moving along microtubules to be transmitted directly to the telomeres (7, 8, 18). We therefore asked whether the pairing defects in *Zcwpw1*^−*/*−^ mice stem from defects in (1) telomere attachment, and/or (2) chromosome movements.

First, by co-staining the shelterin subunit TRF1 with the nuclear lamina marker lamin B1 in WT, *Zcwpw1*^−*/*−^ and *Prdm9*^−*/*−^ testes, we confirmed intact telomere-NE tethering across all genotypes (***SI Appendix*, Fig. S5A**). However, *Zcwpw1*^−*/*−^ spermatocytes showed subtle differences in telomere distribution, including a modest delay in bouquet exit (***SI Appendix*, Fig. S5B**,**C**), which could reflect reduced mobility, a prophase delay (38), or early arrest as observed in a few other meiotic mutants (39–41).

To directly assess chromosome dynamics, we performed live imaging on mouse seminiferous tubule tissue explants stained with Hoechst dye as previously described (18). As expected, WT leptotene/zygotene spermatocytes exhibited prominent RPMs. In sharp contrast, *Zcwpw1*^−*/*−^ leptotene/zygotene spermatocytes showed near complete absence of chromosome movement and nuclear rotation, a phenotype mirroring *Kash5*^−*/*−^ and more severe than *Sun1*^−*/*−^ spermatocytes (18) (**Fig. 4A** and **Supplementary Movies 1–3**). Importantly, RPMs are unaffected in *Prdm9*^−*/*−^ spermatocytes (**Fig. 4A**). Quantitative tracking across WT, *Zcwpw1*^−*/*−^, and *Prdm9*^−*/*−^ confirmed drastic reductions in displacement length (the cumulative distance traveled by tracked chromatin over time in a time-lapse dataset) (WT: 3.6 ± 1.6 μm; *Zcwpw1*^−*/*−^: 0.65 ± 0.71 μm; *Prdm9*^−*/*−^: 3.4 ± 1.5 μm) and speed (WT: 35.0 ± 16.0 nm/s; *Zcwpw1*^−*/*−^: 11.1 ± 9.0 nm/s; *Prdm9*^−*/*−^: 34.3 ± 18.8 nm/s) only in *Zcwpw1*^−*/*−^ (**Fig. 4B,C**). This defect in *Zcwpw1*^−*/*−^ is also different from that previously published in recombination and synapsis mutants, *Dmc1*^−*/*−^ and *Sycp3*^−*/*−^ (18), respectively (reduced but substantial RPMs), suggesting that RPM loss in *Zcwpw1*^−*/*−^ is not due to recombination or synapsis defects alone. Based on these data, we conclude that ZCWPW1 is uniquely required for functional RPMs in a PRDM9-independent manner and that loss of RPMs in *Zcwpw1*^−*/*−^ mice may explain homolog pairing defects, ultimately causing synapsis and DSB repair defects. Together, these data demonstrate that ZCWPW1 acts as an intranuclear regulator of RPMs.

**Figure 4.**
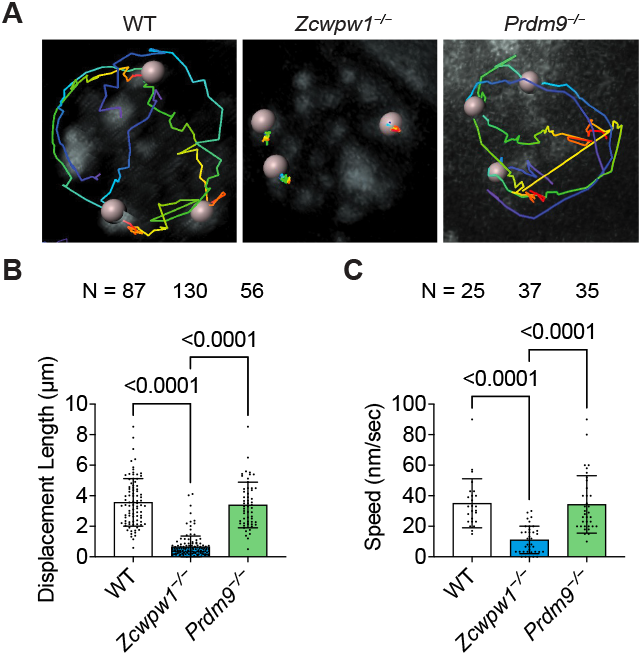
Rapid prophase movements require ZCWPW1 but not PRDM9. (*A*) Trajectory of three heterochromatin spots in early zygotene spermatocyte nuclei of WT, *Zcwpw1*^−*/*−^, and *Prdm9*^−*/*−^ mice. (*B–C*) Quantitation of spot displacement length (*B*) and speed (*C*) in zygotene spermatocytes of WT (n = 3), *Zcwpw1*^−*/*−^ (n = 5), and *Prdm9*^−*/*−^ (n = 4) mice. n: biological replicates. N: spots/blobs quantified per cell are variable, at least 3/per cell. Data shown as mean ± s.d. *P* values were calculated using Kruskal–Wallis tests.

### ZCWPW1 loss in spermatocytes compromises mechanical coupling of telomeres and the cytoskeletal motors

To determine the cause for RPM loss in *Zcwpw1*^−*/*−^ spermatocytes, we next examined the expression and localization of various proteins implicated in telomere attachment or movement. In mammalian meiosis, telomeres are anchored to the nuclear envelope via the shelterin protein (TRF1) (42, 43), which binds the TERB1–TERB2–MAJIN complex (TTM complex) (7, 14, 44) and connects to the LINC complex (composed of inner nuclear membrane SUN proteins (SUN1 (9), SUN2 (45)) and outer nuclear membrane KASH proteins (KASH5 (8, 10))). The LINC complex, in turn, connects telomeres to cytoplasmic dynein-1 (dynein) motors (11) (Fig. 5A).

**Figure 5.**
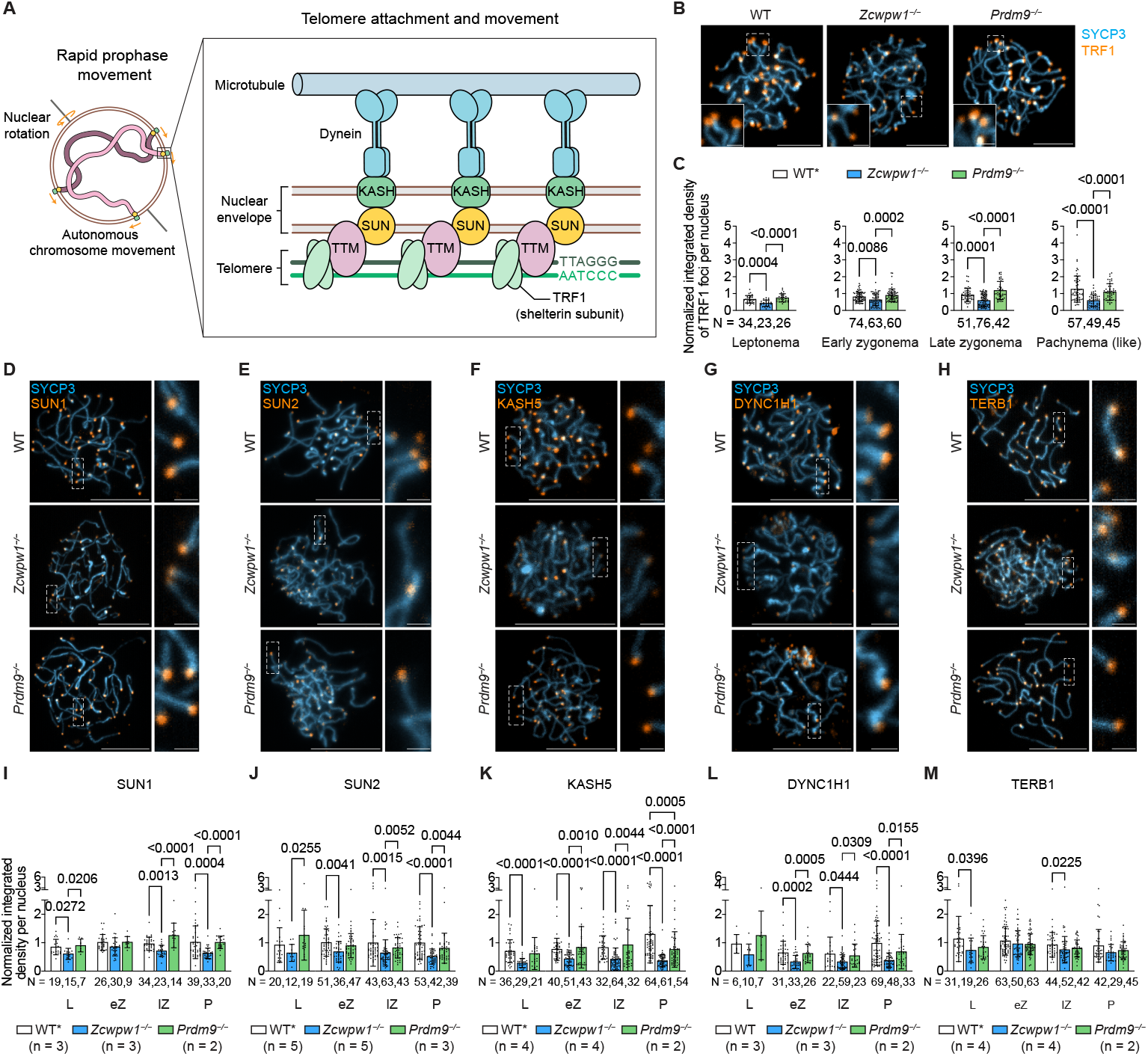
Loss of ZCWPW1 disrupts LINC complex and dynein accumulation at telomeres. (*A*) Schematic of the core molecular machinery for telomere attachment and movement during rapid prophase movement. (*B*) Immunofluorescence of SYCP3 and TRF1 on squashed spermatocytes at pachynema or pachynema-like stages. Scale bars, 5 μm (main images); 1 μm (insets). (*C*) Quantification of the fluorescence intensity of TRF1 during meiotic prophase I of *Zcwpw1*^*+/+*^ or *Zcwpw1*^*+/*−^ (n = 6), *Zcwpw1*^−*/*−^ (n = 6), and *Prdm9*^−*/*−^ (n = 4) mice. Data shown as mean ± s.d. n: biological replicates; N: cells analyzed per group. *P* values were calculated using Ordinary one-way ANOVA (Leptonema) or Kruskal-Wallis tests (other stages). (*D–H*) Immunofluorescence of SYCP3 and SUN1 (*D*), SUN2 (*E*), KASH5 (*F*), DYNC1H1 (*G*), and TERB1 (*H*) on squashed spermatocytes at pachynema or pachynema-like stages. Scale bars, 10 μm (main images); 1 μm (insets). (*I–M*) Quantification of the fluorescence intensity of SUN1 (*I*), SUN2 (*J*), KASH5 (*K*), DYNC1H1 (*L*), and TERB1 (*M*) during meiotic prophase I. Data shown as mean ± s.d. *P* values were calculated using Kruskal–Wallis tests.

Although total cellular protein levels of shelterin and LINC components were not altered in *Zcwpw1*^−*/*−^ (***SI Appendix*, Fig. S5D**), their enrichment at telomeres was significantly reduced (**Fig. 5B–G,I–L**). Quantitative immunofluorescence showed that TRF1 intensity at chromosome ends was maintained in *Prdm9*^−*/*−^ but markedly reduced in *Zcwpw1*^−*/*−^ (WT : *Zcwpw1*^−*/*−^ : *Prdm9*^−*/*−^ mean ratios, 1 : 0.64 : 1.12 (leptonema); 1 : 0.78 : 1.11 (early zygonema); 1 : 0.64 : 1.28 (late zygonema); 1 : 0.46 : 0.86 (pachynema (like)), **Fig. 5B,C**). Furthermore, this reduction in *Zcwpw1*^−*/*−^ telomeres was TRF1-specific, as TRF2 levels were unchanged (***SI Appendix*, Fig. S5E**,**F**). Despite reduced TRF1, TERB1 enrichment was only marginally decreased in *Zcwpw1*^−*/*−^ or *Prdm9*^−*/*−^, suggesting that when TRF1 is reduced, the TTM complex may preserve telomere - nuclear envelope attachment by directly engaging telomeric DNA (46) (**Fig. 5H,M**). In contrast, telomeric enrichment of LINC complex proteins (SUN1, SUN2, KASH5), as well as the dynein heavy chain (DYNC1H1), was significantly reduced in *Zcwpw1*^−*/*−^ spermatocytes (**Fig. 5D–G,I–L**). Taken together, these findings suggest that ZCWPW1 plays a crucial role in the normal recruitment and stability of LINC complex components and dynein. This reduction in dynein may be due either to the loss of KASH5 or loss of intranuclear tension.

### Subtelomeric chromatin structure is altered in *Zcwpw1*^−*/*−^ spermatocytes

The reduction in telomere-bound TRF1 in *Zcwpw1*^−*/*−^ spermatocytes prompted us to investigate potential alterations in the telomere chromatin structure. As a decrease in TRF1 is known to negatively regulate telomere length (47), we measured telomeric repeats using a denatured TeloFISH protocol. Denatured TeloFISH shows that *Zcwpw1*^−*/*−^ telomeres occupied a significantly larger area per nucleus at all substages, but interestingly the integrated telomeric signal was largely comparable to WT with only a modest increase at late zygonema and pachynema. This combination of expanded area without proportional increase in FISH signal intensity suggests altered telomere packaging and early decompaction, with possible lengthening emerging only later in prophase (**Fig. 6A**,**B**). These findings suggested that ZCWPW1 may influence the occupancy or assembly of TRF1 and/or LINC complex directly or indirectly. To probe how ZCWPW1 promotes the stable recruitment or retention of TRF1 and the LINC complex at telomeres, we performed immunoprecipitation followed by mass spectrometry to identify potential interactors. Among this short list, we found several chromatin and telomere/nuclear membrane interactors, including cohesin complex proteins, SUN1, SUN2, lamin B1, and nucleoporins (***SI Appendix*, Table S1**). To determine if these proteins directly interact with ZCWPW1, we ran AlphaFold and identified a strong potential interaction between STAG3 and ZCWPW1 (***SI Appendix*, Fig. S6A**,**B**). STAG3 is a component of the meiosis-specific cohesin that was previously shown to enrich at telomeres and is responsible for protecting telomere integrity and supporting telomere attachment (7). Co-immunoprecipitation in HEK293T cells confirms the interactions between ectopically expressed ZCWPW1 and STAG3 (**Fig. 6D**). Further, immunofluorescence of STAG3 revealed reduced signals near telomeres in *Zcwpw1*^−*/*−^ spermatocytes (**Fig. 6E**). Consistent with reduced cohesin loading, *Zcwpw1*^−/−^ spermatocytes exhibited diminished heterochromatin compaction as indicated by decreased chromocenter intensity **(*SI Appendix*, Fig. S6C**). These chromatin change appear confined to telomere-adjacent regions (subtelomeres and pericentromeres), as the DNA-PAINT signal size of chr.1 and chr.19 doesn’t change relative to the nucleus (***SI Appendix*, Fig. S6D**), nor does the length of the corresponding chromosome axes differ between WT and *Zcwpw1*^−*/*−^ spermatocytes (***SI Appendix*, Fig. S6E**), suggesting that the large-scale chromatin architecture was not substantially affected by the loss of ZCWPW1. Finally, electron microscopy analysis confirms that although telomeres attach in the *Zcwpw1*^−*/*−^ spermatocytes, the telomere attachment plates are weaker, less electron dense, and the surrounding heterochromatin is loosely organized in *Zcwpw1*^−*/*−^ spermatocytes (orange arrowheads, **Fig. 6F**), further underscoring an altered telomere architecture. In summary, loss of ZCWPW1 disrupts telomeric chromatin, which in turn weakens LINC-motor reinforcement at chromosome ends and compromises telomere-led nuclear dynamics.

**Figure 6.**
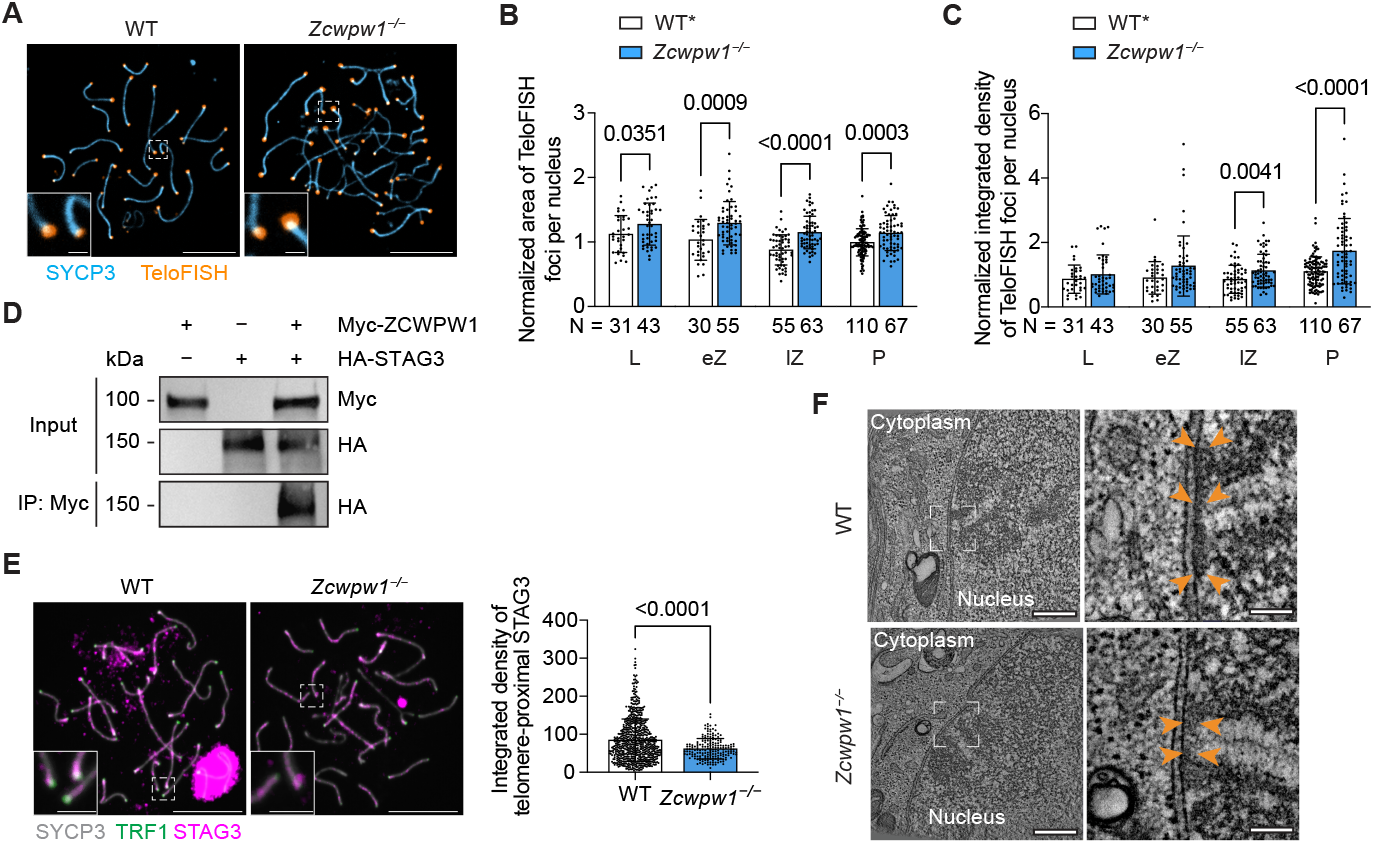
Telomere elongation and decompaction are correlated with reduced cohesin at telomeres in *Zcwpw1*^−*/*−^ spermatocytes. (*A*) Immunofluorescence of SYCP3, and TeloFISH in spermatocytes at pachynema or pachynema-like stages. Scale bars, 5 μm (main images); 1 μm (insets). (*B–C*) Quantification of the area (*B*) and the integrated density (*C*) of TeloFISH (mean ± s.d.; two-tailed Mann–Whitney tests) during meiotic prophase I of WT or *Zcwpw1*^*+/*−^ (n = 4) and *Zcwpw1*^−*/*−^ (n = 4) mice. L: leptonema; eZ: early zygonema; lZ: late zygonema; P: pachynema or pachynema-like. (*D*) Co-immunoprecipitation of ectopically expressed ZCWPW1 and STAG3 in HEK293T cells. (*E*) Left: immunofluorescence of SYCP3, STAG3, and TRF1 on spread spermatocytes from WT and *Zcwpw1*^−*/*−^ mice. Scale bars: 10 μm (main images); 2 μm (insets). Right: quantification of telomere-adjacent STAG3 intensity (mean ± s.d; Mann–Whitney test) in pachynema or pachynema-like cells from WT and *Zcwpw1*^−*/*−^ mice respectively. (*F*) Electron microscopy images of nuclear envelope-attached synaptonemal complex in WT and *Zcwpw1*^−*/*−^ spermatocytes. Telomere attachments were shown in the zoom-in insets. Scale bars: 3 μm (main images); 6 μm (insets).

## Discussion

Taken together, our findings reveal that the loss of ZCWPW1 disrupts the telomere/subtelomere chromatin architecture and the mechanical integrity of meiotic telomeres, ultimately leading to the loss of RPMs. We propose that ZCWPW1 organizes a telomeric hub, maintaining chromatin compaction and mechanical coupling essential for homolog pairing and efficient DSB repair. In *Zcwpw1*^−*/*−^ spermatocytes, telomeres become lengthened and decompacted and have reduced cohesin loading, creating a less rigid anchoring interface with the nuclear envelope. Although attachment is weakly preserved through nearly unaltered TTM complex levels, the concurrent reduction in TRF1, LINC components, and dynein motors weakens the coupling between telomeres and the cytoskeleton motors. This mismatch between a relaxed chromatin interface and reduced motor density dilutes force transmission, rendering the telomeres tethered yet immobile (48) (**Fig. 7**). This phenotype is consistent with simulation studies showing that tethering alone is insufficient to promote pairing and instead slows pairing, whereas the active telomere movements dramatically accelerate pairing (49, 50). Partial detachment, as in *Trf1*^−*/*−^ spermatocytes, permits residual mobility through Brownian diffusion (42), whereas full detachment, as in *Terb1*^−*/*−^ (7), or immobile attachment, as in *Zcwpw1*^−*/*−^, causes pairing failure that triggers meiotic checkpoints and prophase arrest. Therefore, meiosis requires an intricate equilibrium: telomeres must be securely tethered yet dynamically engaged, with the forces they transmit serving as a central determinant of homolog juxtaposition and the fidelity of recombination.

**Figure 7.**
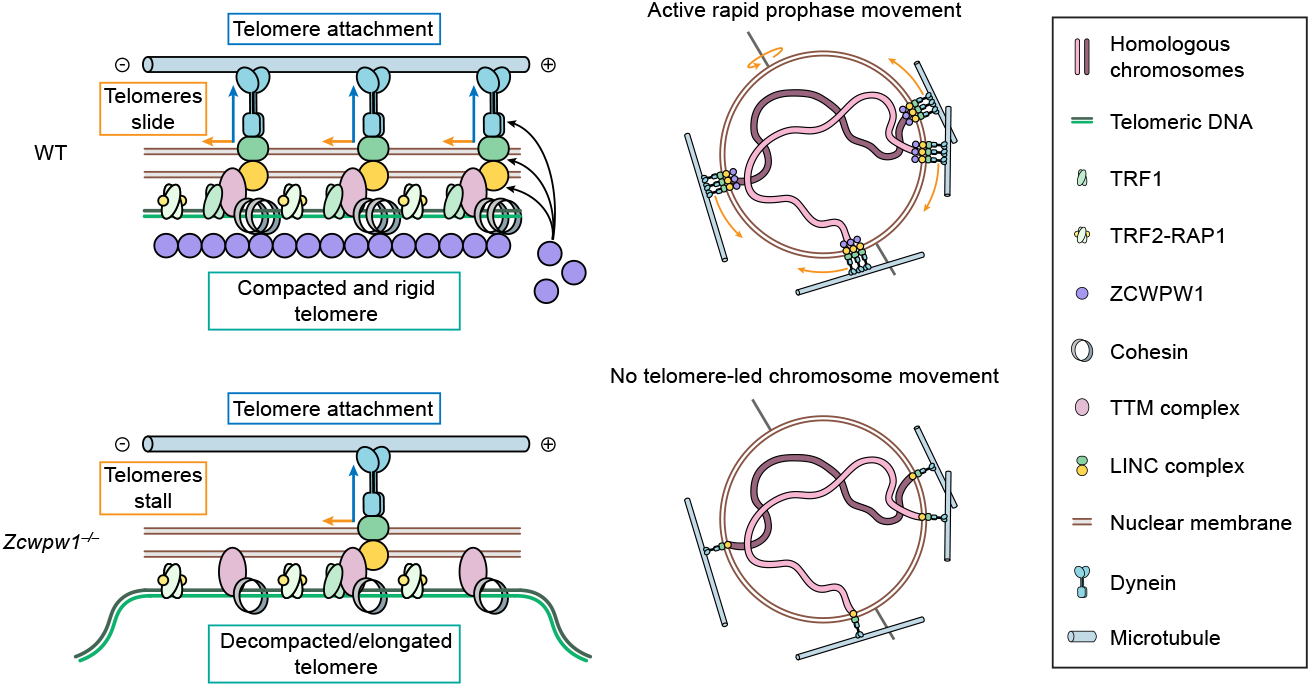
Model of ZCWPW1 in rapid prophase movements (RPMs). ZCWPW1 enriches near telomeres and is responsible for maintaining telomeric cohesin and TRF1 levels. The properly compacted and rigid telomeres and abundant TRF1 allow for stable connection to the nuclear envelope and dynein motor through the LINC complex and efficient force transmission for active RPMs. In *Zcwpw1*^−*/*−^ spermatocytes, telomeres are decompacted and elongated due to loss of TRF1 and telomeric cohesin. The resulting reduction in dynein-generated forces is further compromised by less rigid telomeres, which are unable to overcome friction to drive movements.

Although ZCWPW1 has been viewed as a downstream effector of PRDM9 in DSB repair, due to its PRDM9-dependent histone reading capacity and their co-evolution, our findings uncover an unexpected, PRDM9-independent role in telomere architecture and dynamics. How this functional innovation emerged is unclear, but evolutionary context offers clues. In mammals, *Zcwpw1* has a paralog, *Zcwpw2*, that likely arose through gene duplication. This duplication allowed for functional specialization: ZCWPW2, the shorter paralog containing only chromatin-reading domains, appears to have retained the canonical role in recruiting the DSB machinery, while ZCWPW1 acquired an additional PRDM9-independent role at telomeres. Whether this telomeric function of ZCWPW1’s requires its histone-binding activity is unclear. Notably, knock-in mutations in the H3K4me3-binding domain phenocopy the meiotic defects of *Zcwpw1* deletion, but because RPMs were not assayed in these mutants, we cannot determine whether this role extends to chromosome movement (27). Curiously though human subtelomeres frequently harbor CpG islands and are enriched for H3K4me3, which promotes TERRA (TElomeric Repeat-containing RNA) transcription (51), raising the possibility that chromatin recognition domain guides ZCWPW1 recruitment. While ZCWPW1 preferentially binds dual-modified H3K4me3/H3K36me3 tails (25, 26), it can also localize to sites marked by single H3K4me3 modification in promoter regions as seen in *Prdm9*^−*/*−^ spermatocytes (33). Despite the importance of PRDM9 in recombination, it has been independently lost in several vertebrate lineages including birds, certain reptiles, amphibians, and fishes, while ZCWPW1 is retained (28). Interestingly, telomere-led chromosome movement is highly conserved in eukaryotes. This evolutionary pattern, together with our functional data, suggests that ZCWPW1’s role in telomere architecture and nuclear dynamics reflects an ancestral, PRDM9-independent mechanism that safeguards homolog pairing, even in species lacking PRDM9-directed recombination.

Our work shows that loss of ZCWPW1 leads to chromatin architectural abnormalities observed at telomeric ends. In WT spermatocytes, both telomeric repeat and telomere-bound TRF1/TRF2 signals increase during prophase I (**Fig. 5C,E**, and ***SI Appendix*, Fig. S5G and S6A**), and this coincides with an increase in KASH5 recruitment to telomeres (52). Whether telomeres elongate during this stage has been debated, but our findings, together with recent evidence that telomerase colocalizes with telomeres in mouse and human spermatocytes (53) support the view that telomere extension can occur during prophase I (54–56). In the absence of ZCWPW1, if telomeres are indeed elongated, this likely reflects unrestrained telomerase activity when TRF1-mediated restriction is lost. Intriguingly, TERB1 remains stably associated with telomeres and the TTM complex can preserve nuclear envelope tethering. This can be achieved either because the residual TRF1 is sufficient to mediate TERB1 interaction, and/or the TTM complex associates with telomeres directly through its DNA-binding activity (46). Despite the relatively normal attachment, Z*cwpw1*^−*/*−^ telomeres have reduced levels of cohesin at telomeres. This decrease in cohesin may be a result of altered cohesin turnover, either by directly blocking STAG3 or by interfering with accessory factors that stabilize its residence time. Nevertheless, the combination of elongated and decompacted telomeres and reduced cohesin undermines local chromatin compaction and compromises telomere integrity in *Zcwpw1*^−*/*−^ spermatocytes, leading to impaired recruitment of LINC complex components and dynein motors, which in turn causes defects in RPMs. We were particularly surprised that *Zcwpw1*^−*/*−^ led to a loss of KASH5 and dynein motor accumulation at sites of telomere attachment. This finding may indicate a type of “inside-out” mechanism of telomere-LINC complex-motor protein complex formation, where that tension derived from stable telomere attachment inside the nucleus promotes SUN1-KASH5 recruitment, which in turn promotes dynein interaction.

In short, our findings reveal an unexpected role for ZCWPW1 in coupling chromatin architecture to telomere/chromosome mechanics. Cohesin has been shown to maintain subtelomeric/telomere organization and telomeric integrity (7, 51, 57). In meiosis, cohesin function is diversified through several meiosis-specific complexes containing distinct kleisins (REC8 or RAD21L) and SMC paralogs (SMC1α or SMC1β), yet all of them converge on the requirement for STAG3, which is essential for chromatin loading and stability (58). In spermatocytes, SMC1β has been previously shown to promote chromatin compaction at telomeres and limit TERRA transcription in spermatocytes (59). In its absence, increased TERRA expression and R-loop accumulation compromise heterochromatin domains at chromosomal ends, which likely weakens the TRF1–TTM–LINC linkage, leading to defective attachment and altered telomere length (59– 61). STAG3 stabilizes telomere attachment via its interaction with TERB1 (7), and is further required to maintain the localization and stability of other meiosis-specific cohesin subunits (58), highlighting a potential coupling between the chromatin state and telomere attachment/movement. Here we show that STAG3 interacts with ZCWPW1, and this interaction may stabilize cohesin to preserve telomere architecture. Reduced STAG3 leads to subtelomeric chromatin decompaction, destabilizing shelterin and LINC-associated proteins together with their cytoplasmic dynein motors and ultimately impairing rapid prophase movements. These findings reveal a broader mechanistic principle: chromatin architecture at chromosome ends is actively tuned to sustain the force-bearing attachments and nuclear motions essential for meiotic recombination, homolog pairing, and genome integrity.

Finally, our findings reposition ZCWPW1 from a downstream effector of PRDM9 to a chromatin-based intranuclear regulator of rapid prophase movements with distinct roles in recombination, telomere architecture, and chromosome stability. By constraining telomeric chromatin, maintaining cytoskeletal coupling, and shaping recombination outcomes, ZCWPW1 illustrates how chromatin factors can govern both the organization of chromatin and the mechanical forces that mobilize it within the nucleus. Its loss reveals that meiotic fidelity relies on a finely tuned equilibrium of telomere length, compaction, and mobility—a balance especially critical in spermatocytes. This multifunctionality is reminiscent of CDK2 (62, 63), which coordinates crossover designation and telomere dynamics, underscoring a broader principle in which meiotic proteins achieve robustness by engaging distinct machineries at once. More broadly, ZCWPW1 highlights how evolutionary innovation can couple epigenetic recognition with nuclear force transmission to secure faithful gametogenesis.

## Materials and Methods

*Zcwpw1*^−*/*−^ mice in C57BL/6 and DBA/2J genetic background were generated independently by the University of Michigan Transgenic animal core. Mice were housed and euthanized in accordance with the guidelines of University of Michigan Institutional Committee on Use and Care of Animals and the National Research Council Guide for the Care and Use of Laboratory Animals. Details of spread and squashed spermatocytes preparation, histology, immunofluorescence, spermatogenesis synchronization, antibody production, DNA FISH, electron microscopy, RPM measurement, flow cytometry, recombination assay, immunoblotting, cell culture, co-immunoprecipitation, alphafold prediction, evolutionary analysis, image acquisition, processing, analysis, and statistics are described in SI Appendix.

## Acknowledgments

We thank all members of the Pezza, Cole and Hammoud labs for manuscript discussion and feedback. We thank Drs. Keeney and Petkov for transgenic lines, Drs. Paigen, Handel, and Tóth for custom antibodies. The University of Michigan Transgenic Animal Model Core supported this work. This research was supported by National Institute of Health (NIH) grants GM148028 (S.S.H. & F.C.) 1DP2HD091949-01 (S.S.H.), R01HD104680 01 (S.S.H.), R01HD098129 (F.C.), DP2HD087943 (F.C.), T32 Program 5T32HD079342-10 (D.B.), F31 Fellowship 5F31HD113285-02 (D.B.), CPRIT grant R1213 (F.C.), the Andrew Sabin Family Foundation Fellowship (F.C.), HD110990 (R.P.), and Open Philanthropy Grant 2019-199327 (5384) (S.S.H.).

## References

1. F. Baudat, et al., PRDM9 Is a Major Determinant of Meiotic Recombination Hotspots in Humans and Mice. Science 327, 836–840 (2010).

2. E. D. Parvanov, P. M. Petkov, K. Paigen, Prdm9 Controls Activation of Mammalian Recombination Hotspots. Science 327, 835–835 (2010).

3. S. Myers, et al., Drive Against Hotspot Motifs in Primates Implicates the PRDM9 Gene in Meiotic Recombination. Science 327, 876–879 (2010).

4. D. J. Wynne, O. Rog, P. M. Carlton, A. F. Dernburg, Dynein-dependent processive chromosome motions promote homologous pairing in C. elegans meiosis. J. Cell Biol. 196, 47–64 (2012).

5. A. Penkner, et al., The Nuclear Envelope Protein Matefin/SUN-1 Is Required for Homologous Pairing in C. elegans Meiosis. Dev. Cell 12, 873–885 (2007).

6. A. Sato, et al., Cytoskeletal Forces Span the Nuclear Envelope to Coordinate Meiotic Chromosome Pairing and Synapsis. Cell 139, 907–919 (2009).

7. H. Shibuya, K. Ishiguro, Y. Watanabe, The TRF1-binding protein TERB1 promotes chromosome movement and telomere rigidity in meiosis. Nat. Cell Biol. 16, 145–156 (2014).

8. A. Morimoto, et al., A conserved KASH domain protein associates with telomeres, SUN1, and dynactin during mammalian meiosis. J. Cell Biol. 198, 165–172 (2012).

9. X. Ding, et al., SUN1 Is Required for Telomere Attachment to Nuclear Envelope and Gametogenesis in Mice. Dev. Cell 12, 863–872 (2007).

10. H. F. Horn, et al., A mammalian KASH domain protein coupling meiotic chromosomes to the cytoskeleton. J. Cell Biol. 202, 1023–1039 (2013).

11. R. Agrawal, et al., The KASH5 protein involved in meiotic chromosomal movements is a novel dynein activating adaptor. eLife 11, e78201 (2022).

12. A. Yamamoto, R. R. West, J. R. McIntosh, Y. Hiraoka, A Cytoplasmic Dynein Heavy Chain Is Required for Oscillatory Nuclear Movement of Meiotic Prophase and Efficient Meiotic Recombination in Fission Yeast. J. Cell Biol. 145, 1233–1250 (1999).

13. H. Shibuya, et al., MAJIN Links Telomeric DNA to the Nuclear Membrane by Exchanging Telomere Cap. Cell 163, 1252–1266 (2015).

14. Y. Wang, et al., The meiotic TERB1-TERB2-MAJIN complex tethers telomeres to the nuclear envelope. Nat. Commun. 10, 564 (2019).

15. D. Zickler, N. Kleckner, THE LEPTOTENE-ZYGOTENE TRANSITION OF MEIOSIS. Annu. Rev. Genet. 32, 619–697 (1998).

16. A. Mytlis, K. Levy, Y. M. Elkouby, The many faces of the bouquet centrosome MTOC in meiosis and germ cell development. Curr. Opin. Cell Biol. 81, 102158 (2023).

17. M. N. Conrad, et al., Rapid Telomere Movement in Meiotic Prophase Is Promoted By NDJ1, MPS3, and CSM4 and Is Modulated by Recombination. Cell 133, 1175–1187 (2008).

18. C.-Y. Lee, et al., Mechanism and Regulation of Rapid Telomere Prophase Movements in Mouse Meiotic Chromosomes. Cell Rep. 11, 551–563 (2015).

19. P. Oza, S. L. Jaspersen, A. Miele, J. Dekker, C. L. Peterson, Mechanisms that regulate localization of a DNA double-strand break to the nuclear periphery. Genes Dev. 23, 912–927 (2009).

20. M. Kalocsay, N. J. Hiller, S. Jentsch, Chromosome-wide Rad51 Spreading and SUMO-H2A.Z-Dependent Chromosome Fixation in Response to a Persistent DNA Double-Strand Break. Mol. Cell 33, 335–343 (2009).

21. R. K. Swartz, E. C. Rodriguez, M. C. King, A role for nuclear envelope– bridging complexes in homology-directed repair. Mol. Biol. Cell 25, 2461–2471 (2014).

22. F. Lottersberger, R. A. Karssemeijer, N. Dimitrova, T. de Lange, 53BP1 and the LINC Complex Promote Microtubule-Dependent DSB Mobility and DNA Repair. Cell 163, 880–893 (2015).

23. M. Shokrollahi, et al., DNA double-strand break–capturing nuclear envelope tubules drive DNA repair. Nat. Struct. Mol. Biol. 1–12 (2024). 10.1038/s41594-024-01286-7.

24. S. Qin, J. Min, Structure and function of the nucleosome-binding PWWP domain. Trends Biochem. Sci. 39, 536–547 (2014).

25. M. Mahgoub, et al., Dual histone methyl reader ZCWPW1 facilitates repair of meiotic double strand breaks in male mice. eLife 9, e53360 (2020).

26. D. Wells, et al., ZCWPW1 is recruited to recombination hotspots by PRDM9 and is essential for meiotic double strand break repair. eLife 9, e53392 (2020).

27. T. Huang, et al., The histone modification reader ZCWPW1 links histone methylation to PRDM9-induced double-strand break repair. eLife 9, e53459 (2020).

28. M. I. A. Cavassim, et al., PRDM9 losses in vertebrates are coupled to those of paralogs ZCWPW1 and ZCWPW2. Proc. Natl. Acad. Sci. 119, e2114401119 (2022).

29. M. Li, et al., The histone modification reader ZCWPW1 is required for meiosis prophase I in male but not in female mice. Sci. Adv. 5, eaax1101 (2019).

30. A. G. Hinch, et al., The Configuration of RPA, RAD51, and DMC1 Binding in Meiosis Reveals the Nature of Critical Recombination Intermediates. Mol. Cell 79, 689-701.e10 (2020).

31. F. Cole, S. Keeney, M. Jasin, Comprehensive, Fine-Scale Dissection of Homologous Recombination Outcomes at a Hot Spot in Mouse Meiosis. Mol. Cell 39, 700–710 (2010).

32. M. J. Zelazowski, et al., Age-Dependent Alterations in Meiotic Recombination Cause Chromosome Segregation Errors in Spermatocytes. Cell 171, 601-614.e13 (2017).

33. S. Yuan, et al., The histone modification reader ZCWPW1 promotes double-strand break repair by regulating cross-talk of histone modifications and chromatin accessibility at meiotic hotspots. Genome Biol. 23, 187 (2022).

34. M. Frasca, et al., MutSgamma promotes meiotic recombination and homolog pairing in mouse spermatocytes. Genetics iyaf099 (2025). 10.1093/genetics/iyaf099.

35. K. A. Boateng, M. A. Bellani, I. V. Gregoretti, F. Pratto, R. D. Camerini-Otero, Homologous Pairing Preceding SPO11-Mediated Double-Strand Breaks in Mice. Dev. Cell 24, 196–205 (2013).

36. K. Ishiguro, et al., Meiosis-specific cohesin mediates homolog recognition in mouse spermatocytes. Genes Dev. (2014). 10.1101/gad.237313.113.

37. A. N. Shami, et al., Single-Cell RNA Sequencing of Human, Macaque, and Mouse Testes Uncovers Conserved and Divergent Features of Mammalian Spermatogenesis. Dev. Cell 54, 529-547.e12 (2020).

38. A. Storlazzi, et al., Meiotic double-strand breaks at the interface of chromosome movement, chromosome remodeling, and reductional division. Genes Dev. 17, 2675–2687 (2003).

39. H. Scherthan, et al., Mammalian Meiotic Telomeres: Protein Composition and Redistribution in Relation to Nuclear Pores. Mol. Biol. Cell 11, 4189–4203 (2000).

40. A. J. MacQueen, M. P. Colaiácovo, K. McDonald, A. M. Villeneuve, Synapsis-dependent and -independent mechanisms stabilize homolog pairing during meiotic prophase in C. elegans. Genes Dev. 16, 2428–2442 (2002).

41. E. Trelles-Sticken, J. Loidl, H. Scherthan, Bouquet formation in budding yeast: Initiation of recombination is not required for meiotic telomere clustering. J. Cell Sci. 112, 651–658 (1999).

42. L. Wang, et al., Dual roles of TRF1 in tethering telomeres to the nuclear envelope and protecting them from fusion during meiosis. Cell Death Differ. 25, 1174–1188 (2018).

43. J. Zhang, Z. Tu, Y. Watanabe, H. Shibuya, Distinct TERB1 Domains Regulate Different Protein Interactions in Meiotic Telomere Movement. Cell Rep. 21, 1715–1726 (2017).

44. K. Daniel, et al., Mouse CCDC79 (TERB1) is a meiosis-specific telomere associated protein. BMC Cell Biol. 15, 17 (2014).

45. J. Link, et al., Analysis of Meiosis in SUN1 Deficient Mice Reveals a Distinct Role of SUN2 in Mammalian Meiotic LINC Complex Formation and Function. PLOS Genet. 10, e1004099 (2014).

46. J. M. Dunce, et al., Structural basis of meiotic telomere attachment to the nuclear envelope by MAJIN-TERB2-TERB1. Nat. Commun. 9, 5355 (2018).

47. B. Van Steensel, T. De Lange, Control of telomere length by the human telomeric protein TRF1. Nature 385, 740–743 (1997).

48. W. Xie, M. Gowder, D. Bazzano, M. DeSantis, S. S. Hammoud, Rewiring for movements in meiotic prophase: regulators, roles, and evolutionary pathways. Curr. Opin. Genet. Dev. 93, 102366 (2025).

49. W. F. Marshall, J. C. Fung, Modeling meiotic chromosome pairing: nuclear envelope attachment, telomere-led active random motion, and anomalous diffusion. Phys. Biol. 13, 026003 (2016).

50. W. F. Marshall, J. C. Fung, Modeling meiotic chromosome pairing: a tug of war between telomere forces and a pairing-based Brownian ratchet leads to increased pairing fidelity. Phys. Biol. 16, 046005 (2019).

51. Z. Deng, et al., A role for CTCF and cohesin in subtelomere chromatin organization, TERRA transcription, and telomere end protection. EMBO J. (2012). 10.1038/emboj.2012.266.

52. A. Ma, Y. Yang, L. Cao, L. Chen, J. V. Zhang, FBXO47 regulates centromere pairing as key component of centromeric SCF E3 ligase in mouse spermatocytes. Commun. Biol. 7, 1–14 (2024).

53. R. Reig-Viader, et al., Telomeric Repeat-Containing RNA (TERRA) and Telomerase Are Components of Telomeres During Mammalian Gametogenesis. Biol. Reprod. 90, 103, 1–13 (2014).

54. S. Ozturk, Telomerase Activity and Telomere Length in Male Germ Cells. Biol. Reprod. 92, 53, 1–11 (2015).

55. K. Tanemura, et al., Dynamic rearrangement of telomeres during spermatogenesis in mice. Dev. Biol. 281, 196–207 (2005).

56. T. de Lange, et al., Structure and Variability of Human Chromosome Ends. Mol. Cell. Biol. 10, 518–527 (1990).

57. J. H. I. Haarhuis, et al., The Cohesin Release Factor WAPL Restricts Chromatin Loop Extension. Cell 169, 693-707.e14 (2017).

58. J. Hopkins, et al., Meiosis-Specific Cohesin Component, Stag3 Is Essential for Maintaining Centromere Chromatid Cohesion, and Required for DNA Repair and Synapsis between Homologous Chromosomes. PLOS Genet. 10, e1004413 (2014).

59. U. Biswas, et al., Cohesin SMC1β promotes closed chromatin and controls TERRA expression at spermatocyte telomeres. Life Sci. Alliance 6 (2023).

60. C. Adelfalk, et al., Cohesin SMC1β protects telomeres in meiocytes. J. Cell Biol. 187, 185–199 (2009).

61. E. Revenkova, et al., Cohesin SMC1β is required for meiotic chromosome dynamics, sister chromatid cohesion and DNA recombination. Nat. Cell Biol. 6, 555–562 (2004).

62. A. Viera, et al., CDK2 regulates nuclear envelope protein dynamics and telomere attachment in mouse meiotic prophase. J. Cell Sci. 128, 88–99 (2015).

63. N. Palmer, et al., A novel function for CDK2 activity at meiotic crossover sites. PLOS Biol. 18, e3000903 (2020).

64. J. Lange, et al., The Landscape of Mouse Meiotic Double-Strand Break Formation, Processing, and Repair. Cell 167, 695-708.e16 (2016).

