## Supplemental materials for "ZCWPW1 organizes telomeric architecture to drive meiotic chromosome movements"

### Supporting Information for
ZCWPW1 organizes telomeric architecture to drive meiotic chromosome movements

Wenxin Xie[1], Manjunath Gowder[1], Dominic Bazzano[1], Adrienne Niederriter Shami[1], Aastha Pandey[2], Melissa Frasca[2,3], Lakshmi Paniker[2], Luciana Previato de Almeida[4], Binod Kumar Bharati[5,6], Jing Liang[7], Attila Tóth[8], Morgan DeSantis[9], Jayakrishnan Nandakumar[10], Evgeny Nudler[5,6], Shyamal Mosalaganti[9,11], Roberto Pezza[4,12*], Francesca Cole[2,3*], Saher Sue Hammoud[1,13,14,15*†]

[1]Department of Human Genetics, University of Michigan, Ann Arbor, MI 48109
[2]Department of Epigenetics and Molecular Carcinogenesis, The University of Texas MD Anderson Cancer Center, Houston, TX 77054
[3]The University of Texas MD Anderson Cancer Center UTHealth Houston Graduate School of Biomedical Sciences, Program in Genetics and Epigenetics, Houston, TX 77030
[4]Cell Cycle and Cancer Biology Research Program, Oklahoma Medical Research Foundation, Oklahoma City, OK 73104
[5]Department of Biochemistry and Molecular Pharmacology, New York University Grossman School of Medicine, New York, NY 10016
[6]Howard Hughes Medical Institute, NYU Langone Health, New York, NY 10016
[7]BRCF Microscopy Core, University of Michigan, Ann Arbor, MI 48109
[8]Institute of Physiological Chemistry, Faculty of Medicine at the TU Dresden, Fiedlerstrasse 42, Dresden, Germany 01307
[9]Department of Cell and Developmental Biology, University of Michigan, Ann Arbor, MI 48109
[10]Department of Molecular, Cellular, and Developmental Biology, University of Michigan, Ann Arbor, MI 48109
[11]Life Sciences Institute, University of Michigan, Ann Arbor, MI 48109
[12]Department of Cell Biology, University of Oklahoma Health Science Center, Oklahoma City, OK 73104
[13]Department of Obstetrics and Gynecology, University of Michigan, Ann Arbor, MI 48109
[14]Cellular and Molecular Biology Program, University of Michigan, Ann Arbor, MI 48109
[15]Department of Urology, University of Michigan, Ann Arbor, MI 48109

***Co-corresponding Authors:** Roberto Pezza, Francesca Cole, Saher Sue Hammoud.

**This PDF file includes:**

Supporting text
Figures S1 to S6
Tables S1
Legends for Movies S1 to S3
SI References

**Other supporting materials for this manuscript include the following:**

Movies S1 to S3

#### Supporting Information Text

#### Materials and Methods

##### Mice

All animal experiments were carried out with prior approval of the University of Michigan Institutional Committee on Use and Care of Animals, in accordance with the guidelines established by the National Research Council Guide for the Care and Use of Laboratory Animals. Mice were housed in the University of Michigan animal facility, in an environment controlled for light (12-hour light/dark cycle), temperature (21–23 °C), and seasonal humidity (30%–70%), with minimal disturbances and ad libitum access to food and water. The Zcwpw1-knockout mouse model in C57BL/6 and DBA/2J genetic background was generated independently by the University of Michigan Transgenic animal core. This was achieved by deleting the genomic DNA fragment covering exon 4 using the CRISPR-Cas9-mediated genome editing system with gRNA1 targeting 5'-GTTGACTTAAAACCTGTAGT-3' upstream of exon 4 and gRNA2 targeting 5'-CTGCTGTCTTCCTGAGAAAA-3' downstream of exon. The founders were genotyped by polymerase chain reaction (PCR), followed by DNA sequencing analysis in subsequent generations. Genotyping was performed by PCR amplification of genomic DNA extracted from mouse tails. PCR primers for the Zcwpw1 wild-type and mutant allele were 5'-TGATGAGAGCATTTAGCTAGAAATAATGGGA-3' (forward) and 5'-ACACATAAAATACATTCTCCAAACCTGGG-3' (reverse), yielding a 648-base pair (bp) fragment for wild-type or a 300-bp fragment (varies by founder). Prdm9-knockout mice were described previously and gifted by Petko Petkov at Jax Laboratories (strain B6; 129P2-Prdm9tm1Ymat/J, Stock #010719). Mice were genotyped at the Prdm9 locus using the following primers, and standard cycling conditions: HayComF: GGCATCTCTGATCGCCTAAG, HayWTR: ATATCGGGACCCACCTTCTT, HayMutR: CCGTAATGGGATAGGTCACG.

##### Synchronization of spermatogenesis in juvenile male mice

Synchronization of spermatogenesis was perfomred as previously described (1). 2 days postpartum male pups were pipette-fed with the retinoic acid inhibitor WIN 18,446 resuspended in 1% gum tragacanth (100 µg per 1 g of body weight) for 7 consecutive days. At 9 days postpartum, male pups were injected with 100 µg of retinoic acid dissolved in 10 µL of DMSO. Testes were collected at 23.5 days post injection and the cell stages were validated by immunofluorescence.

##### Protein expression and purification

The full-length mouse Zcwpw1 coding sequence was cloned into the pGEX-6p-1 plasmid backbone with an N-terminal GST tag and transformed into Rosetta DE3 cells. An overnight culture was grown in 50 mL of LB at 37 °C overnight and used to inoculate 2 L of LB broth. The culture was allowed to grow at 37 °C until a log phase OD of 0.4–0.6 (3–4 h). The incubator temperature was lowered to 18 °C, and IPTG was added to a final concentration of 100 µM to induce expression of GST-ZCWPW1. Cells were harvested after an overnight growth by ultracentrifugation, and the cell pellet was resuspended in 40 mL of DTT lysis buffer (24 mM Tris, pH 8.0, 500 mM NaCl, 1 mM DTT, 1 mM PMSF, protease inhibitor). The resuspended pellet was lysed by sonication and clarified by centrifugation at 4 °C for 30 min at 30,000 × g, and the supernatant was filtered through a glass fiber filter. GST-sepharose beads equilibrated in lysis buffer were added to the lysate and allowed to nutate for 2 hours at 4 °C. The bead-bound purified protein lysate was pelleted (2000 × g, 10 min) and washed once with lysis buffer and five times with GST wash buffer (25 mM Tris, pH 8.0, 250 mM NaCl, 1 mM DTT). The bound proteins were eluted using reduced Glutathione Elution Buffer (20 mM reduced L-glutathione, pH 8.0, in GST Wash Buffer) and dialyzed into a buffer containing 25 mM Tris (pH 8.0), 100 mM NaCl, 2 mM DTT for 1 h at 4 °C. Protein concentration was determined by Bradford assay, and GST–ZCWPW1 was cleaved with PreScission protease (10 units/mg). Cleaved products were separated from the GST tag by size exclusion chromatography FPLC, and expression, cleavage, and purity were verified by SDS–PAGE with Coomassie staining.

##### Antibody production

After isolation and purity confirmation by SDS–PAGE, FPLC fractions containing the pure ZCWPW1 were pooled and concentrated. The rabbit and guinea pig polyclonal antibodies against

ZCWPW1 were generated at GeneMed Synthesis via immunization of rabbits and guinea pigs respectively with approximately 2 mg of the full-length ZCWPW1 proteins.

**Antibodies**

The following primary antibodies were used: rabbit anti-ZCWPW1 (this study, 1:1000), guinea pig anti-ZCWPW1 (this study, 1:500), mouse anti-SCP3 (Abcam, ab97672, 1:200 or 1:500), rabbit anti-SCP3 (Abcam, ab15093, 1:200), rabbit anti-SCP1 (Abcam, ab15090, 1:200), rabbit anti-SUN1 (Abcam, ab103021, 1:200), rabbit anti-SUN2 (Abcam, ab124916, 1:200), rabbit anti-TRF2 (Novus, NB110-57130SS, 1:200), rabbit anti-HORMAD1 (Proteintech, 13917-1-AP, 1:200), rabbit anti-Lamin B1 (Abcam, ab16048, 1:500), rabbit anti-Nup153 (Leading Biology, AMR11924N, 1:500), mouse anti-Nup153 (abcam, ab24700, 1:500), rabbit anti-RPA32/RPA2 (Abcam, ab10359, 1:200), rabbit anti-RAD51 (Invitrogen, PA5-27195, 1:200), rabbit anti-KASH5 (Invitrogen, PA5-54116, 1:200), rabbit anti-DYNC1H1 (Bethyl, A304-720A, 1:200), rabbit anti-STAG3 (Invitrogen, PA5-63556, 1:100), mouse anti-TRF1 (Abcam, ab10579, 1:200), rat anti-TRF1 (Abcam, ab192629, 1:200), guinea pig anti-H1t (gift from the Handel Lab, 1:500), guinea pig anti-PRDM9 (gift from the Paigen Lab, 1:500), guinea pig anti-TERB1 (gift from the Toth Lab, 1:40), human anti-CREST (Antibodies Inc., Cat No. 15-234, 1:200 or 1:500), mouse anti-phospho-Histone H2A.X (Ser139) (EMD Millipore, 05-636, 1:1000). The following secondary antibodies were used: goat anti-rat IgG (H+L) highly cross-adsorbed Alexa Fluor™ Plus 594 (Invitrogen, A48264), Alexa Fluor® 488 AffiniPure Donkey Anti-Mouse IgG (H+L) (Jackson ImmunoResearch, 715-545-151), Alexa Fluor® 594 AffiniPure Donkey Anti-Guinea Pig IgG (H+L) (Jackson ImmunoResearch, 706-585-148), Alexa Fluor® 647 AffiniPure Donkey Anti-Rabbit IgG (H+L) (Jackson ImmunoResearch, 711-605-152), Alexa Fluor® 594 AffiniPure Donkey Anti-Rabbit IgG (H+L) (Jackson ImmunoResearch, 711-585-152), Goat anti-Rabbit IgG (H+L) Cross-Adsorbed Secondary Antibody, Alexa Fluor™ 405 (Invitrogen, A-31556), Donkey Anti-Rat IgG (H+L), Highly Cross-Adsorbed, CF®405S (Biotium, 20419). All the secondary antibodies were diluted 1:200 for immunofluorescence of spermatocyte spreads or squashes, or 1:1000 for immunofluorescence of testis cyrosections.

**Spermatocyte spreads and squashes**

Chromosome spreads of prophase I spermatocytes were performed as previously described (2). Spermatocyte squashes were prepared as previously described (3) with modifications. Frozen or freshly isolated testes were detunicated and minced by a razor blade on a glass slide in 2% paraformaldehyde (PFA) in 1× PBS containing 0.15% Triton X-100 for 1 min. Spermatocytes were then squashed by placing a coverslip onto the glass slide. The slides were promptly snap-frozen in liquid nitrogen and retrieved once the bubbling ceased. After thawing the slides and carefully removing the coverslips, the squash preparation was washed 3 × 3 min in 1× PBS with gentle agitation, rinsed in distilled water, and proceeded to immunofluorescence immediately.

**Immunofluorescence of mouse spermatocytes**

Slides of spermatocyte spreads or squashes were heated in antigen retrieval buffer (1 mM EDTA, 0.05% Tween-20, pH 8.0 or Antigen Unmasking Solution, Citrate-Based, pH 6.0 (Vector)) for 5–10 min in a microwave. After cooling down, slides were washed 3 × 5 min with 1× TBS, pH 7.6 with 0.05% Tween-20, and blocked with 1× TBS, pH 7.6, supplemented with 0.05% of Triton X-100, 1% normal goat serum, 3% bovine serum albumin (BSA), and 0.05% sodium azide for 30 min at room temperature. Slides were incubated with primary antibody diluted in 1× TBS, pH 7.6, with 0.05% of Triton X-100, 10% normal goat serum, 3% bovine serum albumin (BSA), and 0.05% sodium azide overnight in a humid chamber at 4 °C. Slides were washed 3 × 5 min with 1× TBS with 0.05% Tween-20, then blocked for 30 min and incubated with secondary antibody and DAPI for 1 h at 37 °C in a humid chamber. Slides were washed 3 × 5 min with 1× TBS with 0.05% Tween-20, then rinsed in distilled water, and mounted before air-drying with Vectashield (Vector Labs). For spermatocyte spreads, after rinsing in H2O, slides were washed with 0.4% Photo-Flo 200 twice, air-dried, and the mounted with Vectashield.

**Immunofluorescence of testis sections**

The dissected testis was denunciated and fixed by 2% PFA overnight at 4 °C, washed 3 × 10 min in PBS and transferred to 30% sucrose in PBS for incubation at 4 °C overnight. The samples were embedded in optimal cutting temperature (OCT) compound and stored at −80 °C. The samples were sectioned for 10 μm using Leica cryostat (Leica CM1950). Immunofluorescence of the cryosections was performed following the user guide of M.O.M.® (Mouse on Mouse) Immunodetection Kit (Vector) with slight adjustments. Slides were dried at room temperature for 10 min, fixed with 4% PFA in 1× PBS, and washed in 3 × 5 min 1× PBS. Tissues were permeabilized with 0.1% Triton X-100 in 1× PBS for 15 min and washed 2 × 5 min in 1× PBS. Slides were boiled in Antigen Unmasking Solution, Citrate-Based, pH 6.0 for 20 min, cooled down, and washed 2 × 5 min in 1× PBS. Slides were incubated in working solution of prepared M.O.M. Mouse IgG Blocking Reagent for 1 h followed by 2 × 2 min washes in 1× PBS. Slides were incubated in working solution of prepared M.O.M. Diluent for 5 min and then in primary antibody diluted in M.O.M. Diluent at overnight in a humid chamber at 4 °C. After washes in 1× PBS with 0.1% Tween-20 for 4 × 15 min, working solution of prepared M.O.M. Biotinylated Anti-Mouse IgG Reagent was applied to sections for 10 min. Slides were washed in 1× PBS for 2 × 2 min, and incubated in streptavidin-based detection system with DAPI for 2 h at room temperature in dark. After washes in 1× PBS with 0.1% Tween-20 for 4 × 15 min, slides were mounted with Vectashield for further analysis.

**Immuno-FISH**

Immunofluorescence of spermatocyte spreads were proceeded as above until the water-rinsing step. Slides were fixed in 3% PFA in 1× PBS for 8 min at room temperature and washed 2 × 2 min with 1× PBS. Slides were dehydrated in 70%, 95%, and 100% of ethanol consecutively for 5 min each, and air-dried completely. Slides were applied with 2× SSC with 70% formamide and denatured at 85 ºC for 10 min. Slides were rehydrated with 2× SSC with 50% formamide for 5 min at room temperature. Slides were heated at 37 ºC for 1 h with hybridization solution (2× SSC with 10% (w/v) dextran sulfate, 126 μg/ml E. coli tRNA, 0.1% BSA, 50% formamide, 0.5 mg/ml salmon sperm DNA, and 1 mM ribonucleoside vanadyl complexes). TelC probes (PNA Bio) diluted in hybridization buffer (1:125) were added to slides and hybridized in a humidified chamber for 2 h at 37 ºC. Slides were washed 2 × 30 min in 2× SSC containing 50% formamide, then washed 2 × 2 min in 1× PBS, rinsed with distilled water, and mounted with Vectashield. For DNA-PAINT, immunofluorescence of spermatocyte spreads was performed and imaged using the same procedures as for immunofluorescence-only slides. After imaging, coverslips of slides were removed, and slides were baked at 65 °C for 1 h. Slides were incubated in 2× SSC with 0.5% TritonX-100 at 37 ºC for 30 min in a water bath. Slides were washed with 2× SSC for 2 min, treated with protease for 5 min at room temperature, and washed with 2× SSC for 5 min at room temperature. Slides were dehydrated in 70%, 80%, and 95% ethanol at room temperature for 2 min each, and air-dried. DNA-PAINT probes (MetaSystems Group) were applied to slides which were then sealed with coverslips. Samples and probes were denatured simultaneously by heating slides at 75 °C for 2 min. Probes on slides hybridized in a humidified chamber at 37 ºC overnight. Slides were soaked into 0.4× SSC (pH 7.0 – 7.5) at 72 °C for 2 min and washed with 2× SSC with 0.05% Tween-20 at room temperature for 30 s. Slides were rinsed in distilled water, air-dried, and mounted with DAPI/Antifade (MetaSystems Group) and stored at 4 °C until imaging.

**Electron microscopy**

Testes were dissected from synchronized mice following euthanasia with $CO_2$ and cervical dislocation. One testis was snap frozen in liquid nitrogen for staging by squash preparations, while the other was processed for fixation. After removal of the tunica albuginea in cold DPBS, seminiferous tubules were transferred to fresh cold DPBS buffer, dissociated by gentle shaking, and kept on ice. Tubules were pre-fixed in freshly prepared Karnovsky's fixative (4% paraformaldehyde, 2.5% glutaraldehyde in 0.2 M cacodylate buffer) for 30 min at room temperature followed by 60 min on ice. Samples were washed five times in 50 mM cacodylate buffer and sent immediately to the electron microscopy core facility (UM). The prefixed tubules were suspended in cell freezing medium (amsbio) and were sandwiched between a 200 μm depth planchette and a flat planchette and then cryo-fixed by Leica EM ICE high-pressure freezer. The frozen samples were stored in liquid nitrogen before they were transferred to Leica EM AFS2 for freeze substitution. The freeze substitution was first processed with 0.5% glutaraldehyde and 0.1% in tannic acid acetone at −90 °C for 96 h followed by acetone wash and 2% osmium tetroxide in acetone. The samples were then kept at −90 °C for 28 h before the temperature gradually increased to −20 °C and kept at −20 °C for 16 h. Samples were then warmed up to 4 °C and washed with acetone and infiltrated with gradient acetone/Epon mixture. After the resin was polymerized in the oven for 48 h, the metal planchettes were removed by liquid nitrogen and hot water. The sample blocks were sent to the electron microscopy core facility at EMBL (Heidelberg, Germany) for sectioning imaging.

**Image acquisition**

Images of spermatocyte spreads and squashes, and OCT-embedded testis sections were acquired on a Nikon A1R-HD25 confocal microscope equipped with a 63× oil-immersion objective (NA 1.4), or on a Zeiss LSM 980 with Airyscan 2. Z-stack images were acquired with a step size of 0.15–0.26 μm on the Nikon confocal microscope, or with a step size of 0.13 μm on the Zeiss microscope.

**Image analysis**

All immunofluorescence and electron microscopy images were processed using FIJI (ImageJ). For signal intensity quantification, regions of interest (ROIs) were manually or semi-automatically defined based on the target signals. Z-stack images were combined into maximum intensity projections prior to analysis. Brightness and contrast were uniformly adjusted across images within the same batch for clarity and visualization, except when adjustments were made solely for visualization purposes without affecting quantitative analyses. The number, area and corresponding mean fluorescence intensity of each ROI were measured, and integrated density was calculated as the product of area and mean intensity. Integrated density values from the same nucleus were summed and normalized to the control group (WT and HET, if applicable), and the normalized values were compared across experimental conditions. For each nucleus, the DNA-PAINT signal area of chr.1 or chr.19 was normalized to the corresponding DAPI-stained nuclear area, while the synaptonemal complex length of chr.1 or chr.19 was normalized to the square root of the same DAPI area. The position of RAD51 foci was measured as the distance from the focus center to the centromere-poximal or centromere-distal TRF1-labeled telomere along the SYCP3 axis and normalized to the total SYCP3 length.

**Measuring RPMs in mouse spermatocytes**

Acquisition and analysis of 3D time-lapse images in the mouse have been described previously (4). Before RPM measurements, seminiferous tubule explants (obtained from 6–8 weeks old mice) were incubated in DMEM media with 1% DMSO–PBS (control) for 30 min at 32 °C before the assay.

**Flow cytometry**

Spermatocytes were isolated as previously described (2). One testis or ~1.5 testes were decapsulated and transferred to 15 mL of Gey's Balanced Salt Solution (GBSS) containing 0.75 mg/mL of collagenase and shaken for 15 min at 33 °C at 500 rpm, occasionally inverted about 10 times (roughly every 5 min). The supernatant was removed using a transfer pipette, being very careful not to remove any seminiferous tubules, after which the tubules were washed with 10 mL of GBSS, again removing the supernatant. For mutant samples, 15 mL of GBSS containing 0.75 mg/mL of trypsin and 25 μL of DNase I (0.6 Kunits/μL) was added. For WT samples, an additional 25 μL of DNase I was added. The solution was shaken for 15 min at 33 °C at 500 rpm and occasionally inverted as before, after which 0.75 mL of Newborn Calf Serum (NCS) was added. The cells were then separated by repeated pipetting for 3 min using a transfer pipette before being filtered using a 70 μm filter top. The solution was spun for 3 min at ~1,800 rpm in benchtop centrifuge, and the supernatant was removed. Next, 25 μL of DNase I was added, and the pellet was resuspended by flicking the tube. The cells were then washed with 10 mL of GBSS containing 2% NCS, followed by an additional 10 μL of DNase I. The sample was spun for 3 min at ~1,800 rpm, the supernatant was removed, and the pellet was resuspended by flicking the tube. For WT samples, 9 mL of GBSS containing 2% NCS and 24 μL of DNase I were added along with 18 μL of Hoechst 33342 (2.5 mg/mL in DMSO, stored at 4 °C). For mutant samples, 3 mL of GBSS containing 2% NCS and 8 μL of DNAse I were added along with 6 μL of Hoechst 33342. The sample was shaken for 45 min at 33 °C at 500 rpm, after which propidium iodide (1 mg/mL, stored at 4 °C) was added (1.8 μL for WT or 0.6 μL for mutant). The sample was filtered again using a 70 μm filter top and transferred to collection tubes. Cells were sorted using either a BD Aria or BD

Fusion flow cytometer with a UV laser (350 nm argon laser). Using blue and red fluorescence from Hoechst 33342, cells were isolated into different meiotic populations as described in the paper. After sorting, a very small portion of cells was counted with a hemocytometer to gauge the number of cells collected. To assess purity, approximately 20,000 cells were immediately washed with 1× PBS and spun for 5 min at 1,500 rpm in a microcentrifuge. The pellet was resuspended in 80 μL of sucrose warmed to 37 °C and left at room temperature for 5 min. Afterward, 20 μL of the sucrose suspension was dropped onto slides already prepped with 65 μL of 1% PFA with 0.1% TritonX-100 (spread across the slides). These slides were placed in a humidifying chamber, which was left closed at room temperature for 2.5 h, then ajar for 30 min, then open for 30 min, after which the slides were rinsed once with water and twice with water containing a 1:250 Photo-Flo 200 solution. The slides were then air-dried and stained with SYCP3, SYCP1, and γH2AX as described in immunofluorescence. Meanwhile, aliquots of sorted cells were pelleted in a 1.5 mL tube at 750 ×g for 5 min, the supernatant removed, and pellets were snap-frozen in an ethanol/dry ice slurry before being transferred to −80 °C for later use.

**Genomic DNA isolation**
The genomic DNA was isolated using the QIAamp DNA Micro kit from Qiagen. The protocol followed was the "Isolation of Genomic DNA from Urine," starting with resuspending the pellet in 300 μL of ATL buffer (preheated at 70 °C to dissolve any precipitates) and adding 20 μL of proteinase K. This solution was mixed by vortexing for 10 s. The mixture was then incubated at 56 °C for 1 h while shaking at 900 rpm. After incubation, the tube was briefly centrifuged to remove any liquid inside the lid, and 300 μL of AL buffer was added along with 50 μL of room temperature 100% ethanol. This was again vortexed for 10 s and briefly centrifuged. The supernatant was then transferred to a column in a 2 mL collection tube and centrifuged at 8,000 rpm for 1 min in a microcentrifuge. The column was placed in a new collection tube, and 500 μL of AW1 buffer was added. This was centrifuged again for 1 min at 8,000 rpm, and the column was placed in a new collection tube. This step was repeated with AW2 buffer, and after each centrifugation, the flow-through was discarded. The sample was centrifuged for 3 min at full speed (14,000 rpm) to dry, and the column was transferred to a clean 1.5 mL tube. Fifty microliters of 5 mM Tris (pH 7.4) were then added, and the sample was incubated at room temperature for 5 min and then centrifuged for 1 min at full speed. The sample was ready for the recombination assay after incubation overnight at 4 °C.

**Determining amplification efficiency of DNA**
Amplification efficiency on isolated DNA from sorted samples was performed as previously described (2, 5). To determine an estimate of the DNA concentration and quality, a nanodrop was used along with running a dilution series on a 1% agarose gel against a 12 ng/μL DNA control. The number of amplifiable DNA molecules/pg was determined using 24–48 PCRs seeded with 2 amplifiable molecules per reaction. Each PCR consisted of 2 nested reactions. This was based on each sample and each hotspot, and the conditions of the PCRs were the same as those used for the recombination assay. Both rounds of PCR contained 1 ASP and 1 universal primer (Supplementary Table 1), and the amplified PCR product was run on a 1% agarose gel. To calculate the amplification adjustment factor, the number of negative wells was counted and input into the equation $-\frac{1}{2}(\ln(\text{total number of negative wells}/\text{total number of wells})$. For reproducibility, the amplification adjustment factor should be between 0.2 and 0.8. If the amplification adjustment factor was over or under this range, the concentration of the DNA is recalculated and the amplification efficiency assay repeated.

**Recombination assay**
The recombination assay was done as previously described (2, 5). The amount of DNA added was calculated based on the amplification efficiency assay. The input DNA was either ~20 or 40 genomes per well, so the recombinant signal was detectable over the nonrecombinant background. The primer annealing temperature was specifically determined for each lot of 2× Q5 master mix and each batch of 11.1× buffer made. For each primary PCR reaction (total volume of 8 μL), DNA was added to 1× buffer containing 2× Q5 master mix from New England Bio Labs and 0.2 μM of

each primer (1 ASP and 1 universal). These primer pairs were used to non-selectively amplify crossovers, noncrossovers, and nonrecombinant DNA. 35 µL of dilution buffer (10 mM Tris–HCl pH 7.5 and 5 µg/mL sonicated salmon sperm DNA) was added to the primary PCR plate. A total of 1.6 µL of this diluted product was then added to the secondary PCR plate. For the secondary plate, the 1× buffer consisted of 11.1× buffer (10×: 450 mM Tris–HCl pH 8.8, 110 mM $(NH_4)_2SO_4$, 45 mM $MgCl_2$, 67 mM beta-mercaptoethanol, 44 µM EDTA, 10 mM each: dATP, dTTP, dGTP, and dCTP, and 1.13 mg/mL non-acetylated BSA), 12.5 mM Tris-base, 0.2 µM of each primer, 0.25 U of Taq, and 0.05 U of Pfu polymerase to a final volume of 30 µL including the DNA input from the primary plate. The PCR program for the primary PCR at A3 was denaturation (98 °C, 1 min for the first denaturation and 20 s for subsequent steps), annealing (30 s at optimized temperature), and extension (72 °C, 1 min per kb). The secondary PCR for *A3* was the same, except the denaturation temperature was 96 °C and the extension temperature was 68 °C. The PCR program for the primary PCR at 59.5 was denaturation (96 °C, 1 min for the first denaturation and 20 s for subsequent steps), annealing (30 s at optimized temperature), and extension (68 °C, 1 min per kb). The secondary PCR for *59.5* was the same, except the extension temperature was 68 °C. For *A3*, the primary PCR had 27 amplification cycles, while for *59.5*, the primary PCR had 26. For both hotspots, the secondary PCR had 36 cycles of amplification. The entire secondary PCR product was then dot blotted to a positively charged nylon membrane in the 96-well format and genotyped by Southern blotting using allele-specific oligos (ASO) probes.

**Testis extraction and western blot**

Adult or juvenile testes were dissected, decapsulated, and homogenized in RIPA buffer (50 mM Tris–HCl pH 7.5, 150 mM NaCl, 1% NP-40, 0.5% sodium deoxycholate, 0.1% SDS) supplemented with protease and phosphatase inhibitors (Sigma). Lysates were incubated on ice for 30 min with occasional vortexing, followed by centrifugation at 14,000 × g for 15 min at 4 °C. Supernatants were collected, and protein concentrations were determined using a BCA assay (Pierce). Equal amounts of protein (30–50 µg) were resolved on 4–20% SDS–PAGE gels and transferred to PVDF membranes (Thermo Scientific). Membranes were blocked with 5% non-fat milk in 1× TBST (TBS + 0.1% Tween-20) for 1 h at room temperature and incubated with primary antibodies overnight at 4 °C. The following primary antibodies were used: anti-ZCWPW1 (this study, 1:500), anti-SUN2 (Abcam, ab124916, 1:500), anti-KASH5 (Invitrogen, PA5-54116, 1:500), anti-TRF1 (Abcam, ab10579, 1:500), anti-TRF2 (Novus, NB110-57130SS, 1:1000), and anti-GAPDH (Proteintech, 10494-1-AP, 1:5000) as a loading control. After washing with 1× TBST for four times, membranes were incubated with HRP-conjugated secondary antibodies for 1 h at room temperature and washed with 1× TBST for four times. Signals were visualized using enhanced chemiluminescence (ECL, Thermo Scientific) and imaged with an iBright Imaging System (Invitrogen).

**Cell culture and co-immunoprecipitation**

HEK293T cells were maintained in Dulbecco's Modified Eagle's Medium (DMEM) supplemented with 10% fetal bovine serum (FBS) at 37 °C in 5% $CO_2$. pCMV-Myc-ZCWPW1 and pCMV-HA-STAG3 were co-transfected into HEK293T cells using lipofectamine 3000. Transfected cells were lysed in Pierce IP lysis buffer (25 mM Tris–HCl, pH 7.4, 150 mM NaCl, 1 mM EDTA, 1% NP-40, and 5% glycerol) supplemented with protease and phosphatase inhibitors (Sigma), and incubated on ice for 60 min. Lysates were clarified by centrifugation at 15,000 × g for 30 min at 4 °C to remove insoluble debris. The supernatants were incubated overnight at 4 °C with anti-Myc or anti-HA antibodies, followed by incubation with protein A/G magnetic beads for 4 h at 4 °C. Beads were washed three times with IP wash buffer (20 mM HEPES–KOH, pH 8.0, 0.5 M NaCl, 1 mM EDTA, 1% NP-40, 5% glycerol, 1 mM DTT), and bound proteins were eluted in 1× SDS loading buffer containing 1% SDS by heating at 95 °C for 10 min. Eluted samples were analyzed by immunoblotting.

**Structural prediction of protein–protein interactions with AlphaFold 3**

Protein–protein interaction models were generated using AlphaFold 3 (6). Full-length amino acid sequences of STAG3 (UniProt: O70576] and residues 241–419 of ZCWPW1 (UniProt: Q6IR42) were used as input. Five models were generated per complex, and the top-ranked prediction based on the interface predicted TM-score (ipTM) and overall confidence metrics was selected for further analysis. Model reliability was assessed using per-residue predicted Local Distance Difference Test (pLDDT) scores, Predicted Aligned Error (PAE) plots, and ipTM values. Structural visualization and interface analysis were performed in UCSF ChimeraX. Only high-confidence interface regions (ipTM > 0.7, pLDDT > 70) were considered for interpretation.

**Protein structure uncertainty and sequence conservation visualization**

Protein domain structure of ZCWPW1 were illustrated using IBS 2.0 (Illustrator for Biological Sequences) (7). Predicted aligned error (PAE) matrices generated by AlphaFold were reduced to two-dimensional representations by averaging across one dimension to obtain residue-level PAE scores. The resulting values were plotted as line graphs using custom Python scripts (NumPy and Matplotlib). For sequence conservation analysis, homologous protein sequences with at least 50% identity were retrieved from UniProt, and multiple sequence alignments were generated. Residue-level Jensen–Shannon divergence (JSD) scores were then calculated from these alignments and plotted with Python.

**Statistical analysis and reproducibility**

All experiments were performed with at least two independent biological replicates; the exact sample sizes are provided in the corresponding figure or figure legends. Statistical analyses and graph generation were conducted using GraphPad Prism 10 or Python 3. Normality of each dataset was assessed using the Kolmogorov–Smirnov, Anderson–Darling, D'Agostino–Pearson omnibus, or Shapiro–Wilk test, depending on sample size and distribution characteristics. For datasets with small sample sizes, normal Q–Q plots were also examined to assess normality. For data following a normal distribution with similar standard deviations across groups, two-tailed unpaired t-tests (for two groups) or one-way ANOVA (for three or more groups) were used. For data with unequal variances, Welch's t-test (for two groups) or Welch or Brown–Forsythe ANOVA test (for three or more groups) was applied instead. Non-normally distributed data were analyzed using two-tailed Mann–Whitney U-tests (for two groups) or Kruskal–Wallis tests (for three or more groups). Mixed-effects analysis was used to compare the frequencies of heterologous synapsis and asynapsis across genotypes, treating genotype and phenotype type as fixed effects and individual animals as random effects to account for repeated measures. 2-tailed Chi-square with Yates correction was used to compare the number of noncrossovers. Normalized RAD51 positions (ratio of distance to SC length) were classified into telomeric (0–0.04 and 0.96–1.0), subtelomeric (0.04–0.10 and 0.90–0.96), and central (0.10–0.90) regions. Distributions of RAD51 counts across these regions were compared between genotypes using Chi-square tests. Region-specific enrichment or depletion was further evaluated with Fisher's exact test (region vs. rest), and odds ratios were reported to indicate the direction and magnitude of enrichment. A p-value < 0.05 was considered statistically significant and indicated in figures. Corrections for multiple comparisons and exact p-values are reported in the respective figures. Sample sizes were determined based on prior experimental experience to ensure adequate statistical power. Investigators were not blinded during data collection and quantification, as the phenotypic differences between experimental conditions were visually apparent. Cells were selected at random for analysis.

### Figures

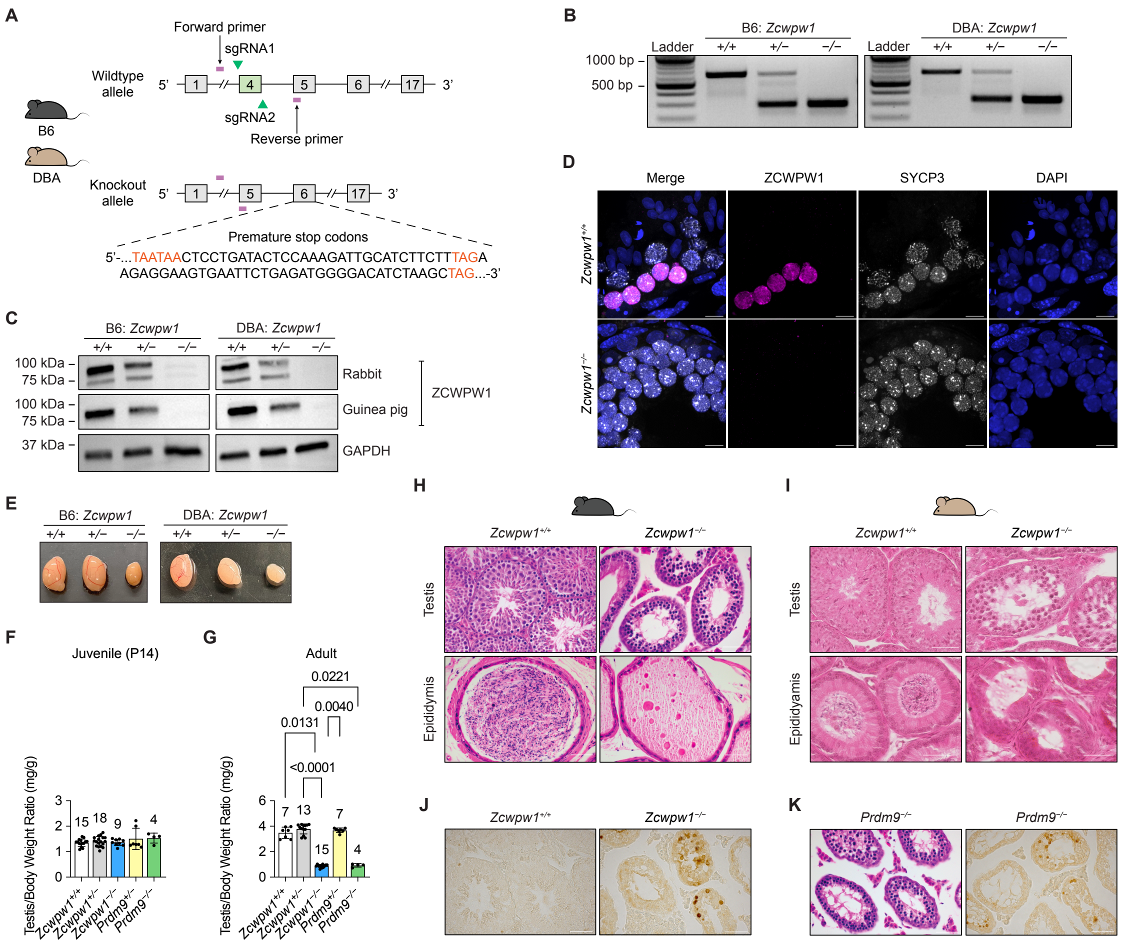


**Fig. S1. Loss of ZCWPW1 results in germ cell deaths, meiotic arrest, and infertility of male mice.** (*A*) Schematics of mouse Zcwpw1 gene structure and targeting strategy used to generate *Zcwpw1*$^{-/-}$ mice. Excision of exon 4 by CRISPR-Cas9 leads to multiple premature termination codons occurring in exon 6. Primers used for genotyping are indicated. (*B*) Genotyping analysis confirms successful gene deletion. (*C*) Western blot demonstrating loss of ZCWPW1 expression in knockout mouse testes. Loading control: GAPDH. (*D*) Immunofluorescence of ZCWPW1 and SYCP3 in representative zygotene cells on testis cross-sections from *Zcwpw1*$^{+/+}$ and *Zcwpw1*$^{-/-}$ B6 mice. Nuclei were stained by 4′,6-diamidino-2-phenylindole (DAPI). Scale bars: 10 μm. (*E*) Representative photo of testes from B6 adult mice (left), and *Zcwpw1*$^{+/+}$ (adult), *Zcwpw1*$^{+/-}$ (6 weeks), and *Zcwpw1*$^{-/-}$ (6 weeks) DBA mice. (*F–G*) Testis (average weight of two testes) to body weight ratio (arbitrary units) for juvenile (*F*) and adult (*G*) B6 mice. Data shown as mean ± s.d. P values were calculated using Kruskal–Wallis test. The number of independent biological replicates are indicated above bars of each corrresponding genotypes. (*H–I*) Testes (left) and epididymis (right) histology of *Zcwpw1*$^{+/+}$ and *Zcwpw1*$^{-/-}$ B6 (adult) (*H*) and DBA (6 weeks) (*I*) mice, as stained with hematoxylin and eosin (H&E). Scale bars: 50 μm. (*J*) Terminal deoxynucleotidyl transferase dUTP nick end labeling (TUNEL) staining of adult testes identify a significant number of apoptotic cells in *Zcwpw1*$^{-/-}$ B6 mice. Scale bars: 50 μm. (*K*) H&E (left) and TUNEL (right) staining in *Prdm9*$^{-/-}$ mouse testes demonstrate a similar spermatogenetic phenotype. Scale bars: 50 μm.

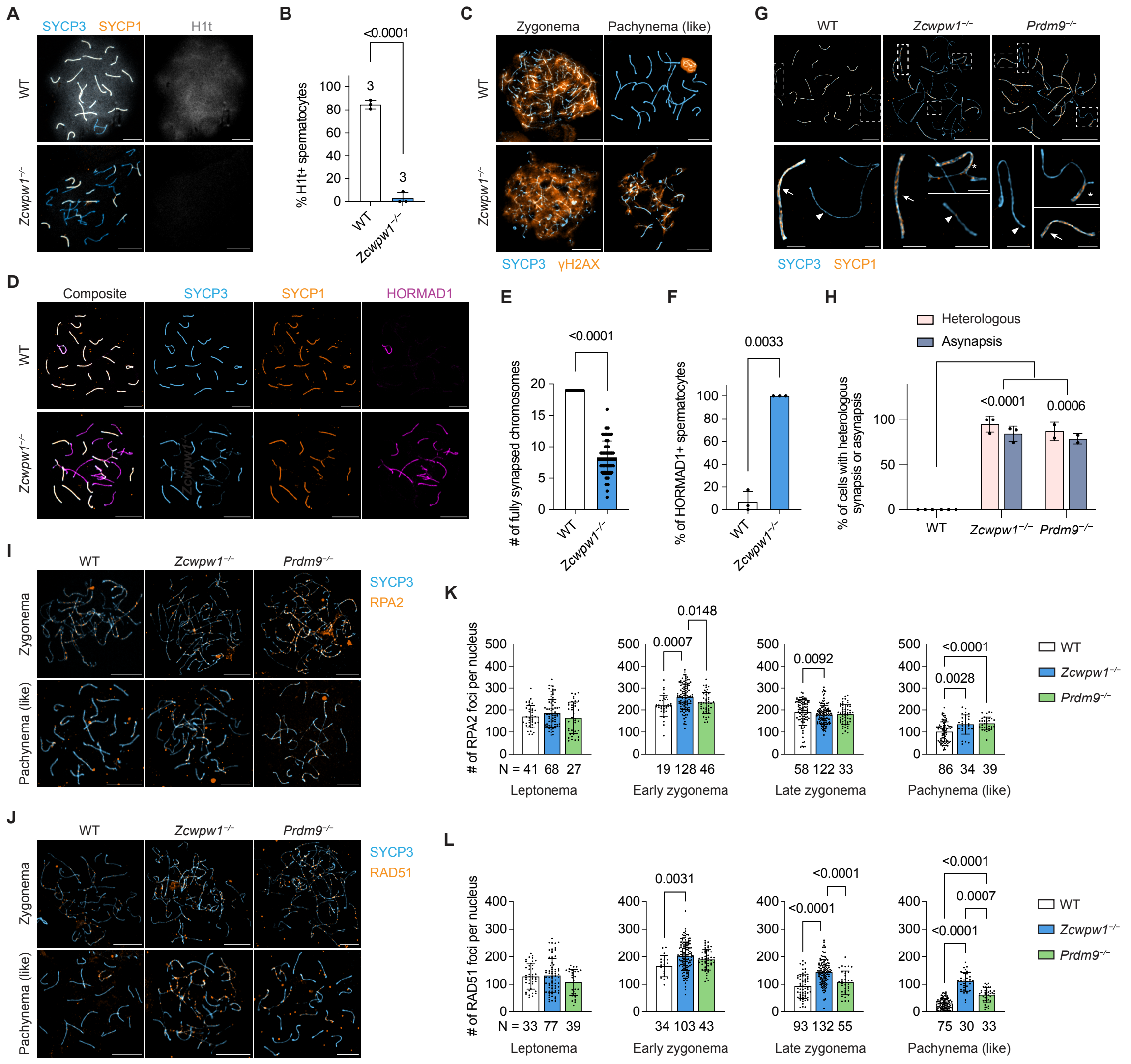


**Fig. S2. *Zcwpw1*$^{-/-}$ spermatocytes are arrested before mid-pachynema with more severe synapsis and DSB repair defects than *Prdm9*$^{-/-}$.** (*A*) Immunofluorescence of SYCP1, SYCP3, and H1t in spread spermatocytes of WT and *Zcwpw1*$^{-/-}$ mice. Scale bars, 10 µm. (*B*) Quantification of cells with H1t signal at pachynema or pachynema-like stages of WT (n = 3) and *Zcwpw1*$^{-/-}$ (n = 3) mice. Data shown as mean ± s.d. *P* values was calculated using Welch's t test. (*C*) Immunofluorescence of SYCP3 and γH2AX in spread spermatocytes. Scale bars, 10 µm. (*D–F*) Immunofluorescence of SYCP1, SYCP3, and HORMAD1 (*D*), quantification of number of fully synapsed chromosomes (*E*), and quantification of HORMAD1 positive (excluding sex chromosomes) (*F*) spermatocytes at pachynema or pachynema-like stage of WT (n = 3) and *Zcwpw1*$^{-/-}$ (n = 3) mice. Scale bars, 10 µm. Data shown as mean ± s.d. *P* value was calculated using Welch's t tests. (*G*) Representative immunofluorescence images of synapsis in spread spermatocytes at pachynema or pachynema-like stages. Different types of synapsis highlighted with dashed squares are shown in the insets. Arrows: homologous synapsis. Arrowheads: asynapsis. Asterisks: heterologous synapsis. Scale bars, 10 µm (main images); 2 µm (insets). (*H*) Quantification of abnormal synapsis in zygonema and pachynema/pachynema-like of WT (n = 3), *Zcwpw1*$^{-/-}$ (n = 3), and *Prdm9*$^{-/-}$ (n = 2) mice. Data shown as mean ± s.d. *P* values were calculated using mixed-effects analysis. (*I–J*) Immunofluorescence of SYCP3 and RPA2 (*I*) or RAD51 (*J*) in spread spermatocytes at zygonema and pachynema or pachynema-like stages. Scale bars, 10 µm. (K–L) Quantification of the number of RPA2 (*K*) or RAD51 (L) foci per nucleus in meiotic prophase

I of WT (n = 3), *Zcwpw1*$^{-/-}$ (n = 3), and *Prdm9*$^{-/-}$ (n = 3) mice. N: number of nuclei analyzed per group. Data shown as mean ± s.d. *P* values were calculated using Welch and Brown–Forsythe ANOVA tests (leptonema and pachynema in *K*; early zygonema in *L*), Ordinary one-way ANOVA (early zygonema in *K*; late zygonema in *L*), and Kruskal–Wallis test (late zygonema in *K*; leptonema, pachynema in *L*).

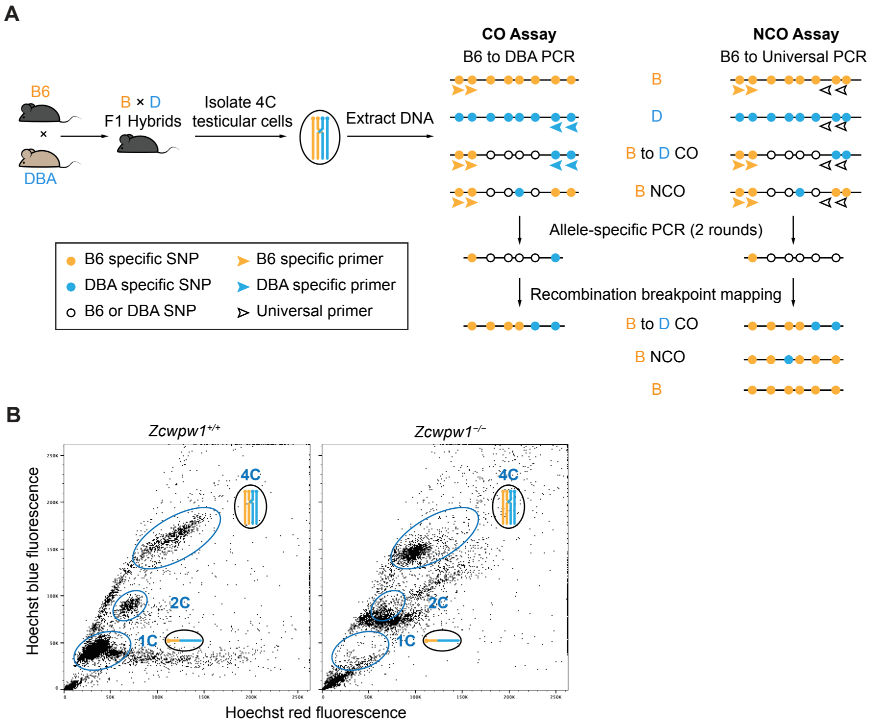


**Fig. S3. Recombination assay workflow.** (*A*) Schematics of recombination assays for detecting COs and NCOs from B6 × DBA F1 hybrid male mice. (*B*) Representative fluorescence-activated cell sorting (FACS) profiles of testicular cells from *Zcwpw1*$^{+/+}$ and *Zcwpw1*$^{-/-}$ mice.

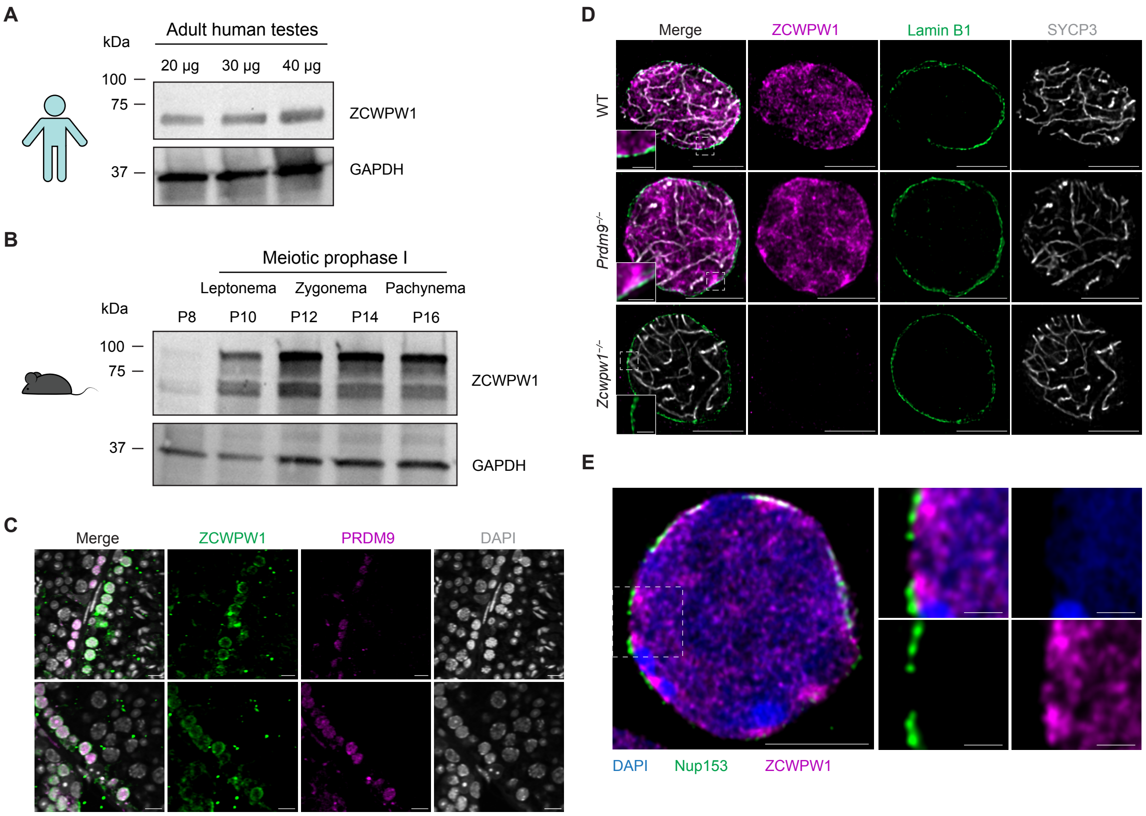


**Fig. S4. ZCWPW1 is expressed in mouse and in human testes, and PRDM9 is not enriched at the nuclear periphery.** (*A*) Western blot of ZCWPW1 protein expression in adult human testes. (*B*) Western blot of ZCWPW1 protein expression during the meiotic prophase I of the first wave of spermatogenesis in mice. (*C*) Immunofluorescence of ZCWPW1 and PRDM9 on FFPE testis cross-sections. Nuclei were stained by 4′,6-diamidino-2-phenylindole (DAPI). Scale bars, 10 µm. (*D*) Representative immunofluorescence images of ZCWPW1 localization in WT, *Zcwpw1*$^{-/-}$, and *Prdm9*$^{-/-}$ mice. Scale bars: 5 µm (main); 1 µm (insets). (*E*) Preferential clustering of ZCWPW1 underneath nuclear pore complexes (NPCs) and away from DAPI-dense regions. Scale bars: 5 µm (main); 1 µm (insets).

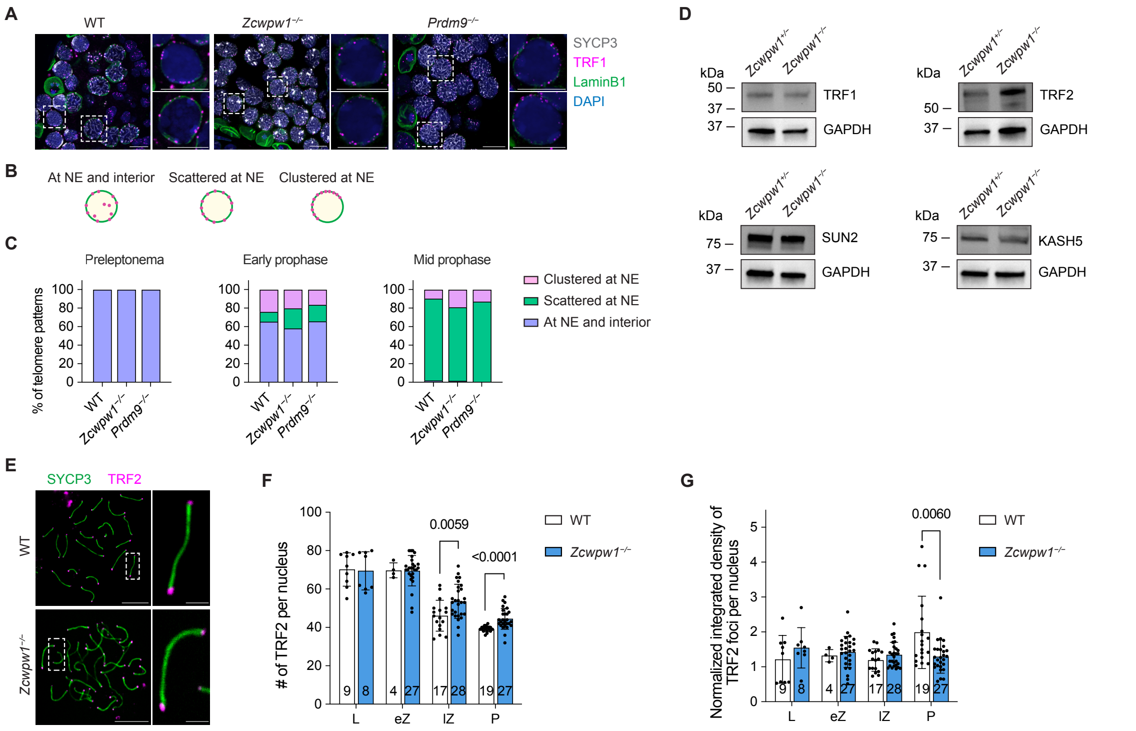


**Fig. S5. Loss of ZCWPW1 does not affect telomere attachment to the nuclear envelope or the total abundance of telomere-associated proteins.** (*A*) Immunofluoresence of SYCP3, lamin B1, and TRF1 on testis cross sections. Merged channels of TRF1, lamin B1, and DAPI of spermatocytes highlighted with dashed squares are shown in the insets. Scale bars, 10 µm. Brightness and contrast were set differently for better visualization. (*B*) Schematics of telomere localization patterns during meiotic prophase I. (*C*) Quantification of different telomere localization patterns across stages in WT (n = 2), *Zcwpw1*$^{-/-}$ (n = 2), and *Prdm9*$^{-/-}$ (n = 1) spermatocytes. (*D*) Western blot of the protein expression of TRF1, TRF2, SUN2, and KASH5 in testes from P14 WT and *Zcwpw1*$^{-/-}$ mice. (*E*) Immunofluorescence of SYCP1, SYCP3, and TRF2 on spread spermatocytes at pachynema or pachynema-like stages. Synaptonemal complexes with corresponding TRF2 foci highlighted with dashed rectangles are shown in insets. Scale bars, 10 µm (main images); 2 µm (insets). (*F*) Quantification of the number of TRF2 foci per nucleus during meiotic prophase I of WT (n = 3) and *Zcwpw1*$^{-/-}$ (n = 3) mice. Cells analyzed per group were shown at the bottom of each bar. Data shown as mean ± s.d. *P* values were calculated using Mann–Whitney tests (L, eZ) and Welch's t tests (lZ, P). (*G*) Quantification of the intensity of TRF2 foci per nucleus during meiotic prophase I of WT (n = 3) and *Zcwpw1*$^{-/-}$ (n = 3) mice. Data shown as mean ± s.d. Cells analyzed per group were shown at the bottom of each bar. *P* values were calculated using Welch's t test (L) and Mann–Whitney tests (eZ, lZ, P).

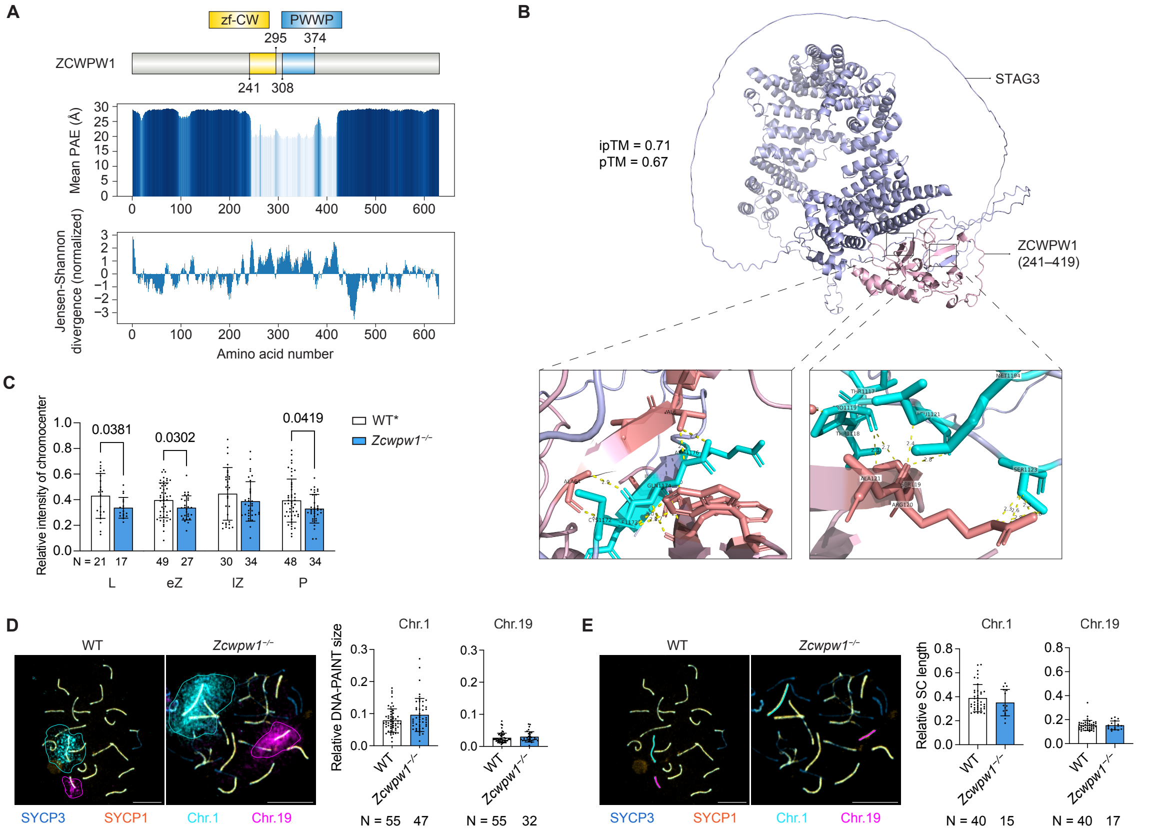


**Fig. S6. Chromatin near telomeres is decompacted in *Zcwpw1*$^{-/-}$ spermatocytes.** (*A*) ZCWPW1 sequence and domain layout are shown (top), the AlphaFold PAE (Predicted Aligned Error) plot highlights structural confidence (middle), and per-residue Jensen–Shannon divergence indicates conservation across homologs with 50% of identity (bottom). (*B*) AlphaFold prediction of the interaction between residues 241–419 of ZCWPW1 and STAG3. Selected residues from ZCWPW1 (salmon sticks) and STAG3 (cyan sticks) are displayed in stick representation. Intermolecular contacts within 3.0 Å are highlighted as yellow dashed lines. (*C*) Quantification of the relative intensity of DAPI-dense regions (chromocenters) per total nucleus from WT or *Zcwpw1*$^{+/-}$ (n = 3) and *Zcwpw1*$^{-/-}$ (n = 3) mice. Cells analyzed per group were shown at the bottom of each bar. Data shown as mean ± s.d. *P* values were calculated using Welch's t test. (*D*–*E*) Representative images (upper) and quantification (lower) of normalized DNA-PAINT size (*D*) of paired chr.1 and chr.19 homologs, and normalized lengths of the synaptonemal complex (*E*) of synapsed chr.1 and chr.19 homologs from WT (n = 4) and *Zcwpw1*$^{-/-}$ (n = 3) mice. Cells analyzed per group were shown at the bottom of each bar. Data shown as mean ± s.d. *P* values were calculated using Mann–Whitney tests. Scale bars: 10 μm.

#### Tables

**Table S1.** Select list of ZCWPW1 interactors identified by IP-MS

| Function | Description | Sequence length | PSM Count | Coverage | PSM count /Sequence Length | NSAF (Normalized Spectral Abundance Factor) | NSAF (%) |
|---|---|---|---|---|---|---|---|
| | Zinc finger CW-type PWWP domain protein 1 OS=Mus musculus OX=10090 GN=Zcwpw1 PE=1 SV=2 | 630 | 18 | 0.165 | 0.029 | 0.044 | 100.0 |
| DNA repair | Serine/threonine-protein kinase ATR OS=Mus musculus OX=10090 GN=Atr PE=1 SV=2 | 2635 | 51 | 0.167 | 0.019 | 0.030 | 67.7 |
| DNA repair | Chromodomain-helicase-DNA-binding protein 4 OS=Mus musculus OX=10090 GN=Chd4 PE=1 SV=1 | 1915 | 34 | 0.124 | 0.018 | 0.028 | 62.1 |
| DNA repair | DNA damage-binding protein 1 OS=Mus musculus OX=10090 GN=Ddb1 PE=1 SV=2 | 1140 | 20 | 0.135 | 0.018 | 0.027 | 61.4 |
| DNA repair | DNA ligase 3 OS=Mus musculus OX=10090 GN=Lig3 PE=1 SV=2 | 1015 | 17 | 0.137 | 0.017 | 0.026 | 58.6 |
| DNA repair | General transcription and DNA repair factor IIH helicase subunit XPB OS=Mus musculus OX=10090 GN=Ercc3 PE=2 SV=1 | 783 | 25 | 0.24 | 0.032 | 0.050 | 111.7 |
| DNA repair | DNA repair protein XRCC1 OS=Mus musculus OX=10090 GN=Xrcc1 PE=1 SV=2 | 631 | 13 | 0.049 | 0.021 | 0.032 | 72.1 |
| DNA repair | Mediator of DNA damage checkpoint protein 1 OS=Mus musculus OX=10090 GN=Mdc1 PE=1 SV=1 | 1707 | 13 | 0.06 | 0.008 | 0.012 | 26.7 |
| DNA repair | DNA damage-binding protein 1 OS=Mus musculus OX=10090 GN=Ddb1 PE=1 SV=2 | 1140 | 13 | 0.045 | 0.011 | 0.018 | 39.9 |
| DNA repair | X-ray repair cross-complementing protein 5 OS=Mus musculus OX=10090 GN=Xrcc5 PE=1 SV=4 | 732 | 9 | 0.036 | 0.012 | 0.019 | 43.0 |
| Nuclear pore complex | Nuclear pore complex protein Nup160 OS=Mus musculus OX=10090 GN=Nup160 PE=1 SV=2 | 1402 | 42 | 0.262 | 0.030 | 0.047 | 104.9 |

| | | | | | | | |
|---|---|---|---|---|---|---|---|
| Nuclear pore complex | Nuclear pore complex protein Nup155 OS=Mus musculus OX=10090 GN=Nup155 PE=1 SV=1 | 1391 | 34 | 0.197 | 0.024 | 0.038 | 85.5 |
| Nuclear pore complex | Nuclear pore complex protein Nup98-Nup96 OS=Mus musculus OX=10090 GN=Nup98 PE=1 SV=2 | 1816 | 31 | 0.134 | 0.017 | 0.027 | 59.7 |
| Chromosome axis and synapsis | Synaptonemal complex protein 2 OS=Mus musculus OX=10090 GN=Sycp2 PE=1 SV=2 | 1500 | 22 | 0.139 | 0.015 | 0.023 | 51.3 |
| Chromosome axis and synapsis | Structural maintenance of chromosomes protein 3 OS=Mus musculus OX=10090 GN=Smc3 PE=1 SV=2 | 1217 | 22 | 0.106 | 0.018 | 0.028 | 63.3 |
| Chromosome axis and synapsis | Structural maintenance of chromosomes protein 6 OS=Mus musculus OX=10090 GN=Smc6 PE=1 SV=1 | 1097 | 22 | 0.172 | 0.020 | 0.031 | 70.2 |
| Chromosome axis and synapsis | Sister chromatid cohesion protein PDS5 homolog B OS=Mus musculus OX=10090 GN=Pds5b PE=1 SV=1 | 1446 | 18 | 0.093 | 0.012 | 0.019 | 43.6 |
| Chromosome axis and synapsis | Structural maintenance of chromosomes protein 5 OS=Mus musculus OX=10090 GN=Smc5 PE=1 SV=1 | 1101 | 18 | 0.065 | 0.016 | 0.025 | 57.2 |
| Chromosome axis and synapsis | Sister chromatid cohesion protein PDS5 homolog A OS=Mus musculus OX=10090 GN=Pds5a PE=1 SV=3 | 1332 | 14 | 0.064 | 0.011 | 0.016 | 36.8 |
| Chromosome axis and synapsis | Synaptonemal complex central element protein 1 OS=Mus musculus OX=10090 GN=Syce1 PE=1 SV=1 | 329 | 10 | 0.112 | 0.030 | 0.047 | 106.4 |

| | | | | | | | |
|---|---|---|---|---|---|---|---|
| Chromosome axis and synapsis | Structural maintenance of chromosomes protein 1B OS=Mus musculus OX=10090 GN=Smc1b PE=1 SV=1 | 1248 | 9 | 0.072 | 0.007 | 0.011 | 25.2 |
| Chromosome axis and synapsis | Synaptonemal complex protein 1 OS=Mus musculus OX=10090 GN=Sycp1 PE=1 SV=2 | 993 | 6 | 0.057 | 0.006 | 0.009 | 21.1 |
| Chromosome axis and synapsis | Cohesin subunit SA-3 OS=Mus musculus OX=10090 GN=Stag3 PE=1 SV=2 | 1240 | 21 | 0.11 | 0.017 | 0.026 | 59.3 |
| Telomere associated | Telomere-associated protein RIF1 OS=Mus musculus OX=10090 GN=Rif1 PE=1 SV=2 | 2419 | 19 | 0.06 | 0.008 | 0.012 | 27.5 |
| Telomere associated | SUN domain-containing protein 2 OS=Mus musculus OX=10090 GN=Sun2 PE=1 SV=3 | 731 | 13 | 0.103 | 0.018 | 0.028 | 62.2 |
| Telomere associated | SUN domain-containing protein 1 OS=Mus musculus OX=10090 GN=Sun1 PE=1 SV=2 | 913 | 5 | 0.046 | 0.005 | 0.009 | 19.2 |
| Telomere associated | Lamin-B1 OS=Mus musculus OX=10090 GN=Lmnb1 PE=1 SV=3 | 588 | 15 | 0.088 | 0.026 | 0.040 | 89.3 |
| Telomere associated | Lamin-B2 OS=Mus musculus OX=10090 GN=Lmnb2 PE=1 SV=2 | 596 | 8 | 0.067 | 0.013 | 0.021 | 47.0 |
| Chromatin remodeller | SWI/SNF complex subunit SMARCC2 OS=Mus musculus OX=10090 GN=Smarcc2 PE=1 SV=2 | 1213 | 15 | 0.106 | 0.012 | 0.019 | 43.3 |
| Chromatin remodeller | Histone deacetylase 1 OS=Mus musculus OX=10090 GN=Hdac1 PE=1 SV=1 | 482 | 11 | 0.137 | 0.023 | 0.035 | 79.9 |
| Chromatin remodeller | Histone deacetylase 2 OS=Mus musculus OX=10090 GN=Hdac2 PE=1 SV=1 | 488 | 9 | 0.117 | 0.018 | 0.029 | 64.5 |
| Chromatin remodeller | Histone deacetylase 6 (Fragment) OS=Mus musculus OX=10090 GN=Hdac6 PE=1 SV=1 | 1008 | 6 | 0.079 | 0.006 | 0.009 | 20.8 |

| | | | | | | | |
|---|---|---|---|---|---|---|---|
| Topoisomerase | DNA topoisomerase 2-alpha OS=Mus musculus OX=10090 GN=Top2a PE=1 SV=2 | 1528 | 49 | 0.135 | 0.032 | 0.050 | 112.2 |
| Topoisomerase | DNA topoisomerase 2-beta OS=Mus musculus OX=10090 GN=Top2b PE=1 SV=2 | 1612 | 41 | 0.117 | 0.025 | 0.039 | 89.0 |
| Topoisomerase | DNA topoisomerase 1 OS=Mus musculus OX=10090 GN=Top1 PE=1 SV=2 | 767 | 13 | 0.059 | 0.017 | 0.026 | 59.3 |

**Movie S1. Active rapid prophase movements in WT nuclei.**

**Movie S2. Rapid prophase movements are lost in *Zcwpw1*$^{-/-}$ nuclei.**

**Movie S3. Rapid prophase movements are preserved in *Prdm9*$^{-/-}$ nuclei.**

#### SI References


1. C. A. Hogarth, *et al.*, Turning a Spermatogenic Wave into a Tsunami: Synchronizing Murine Spermatogenesis Using WIN 18,4461. *Biol. Reprod.* **88**, 40, 1–9 (2013).

2. M. J. Zelazowski, *et al.*, Age-Dependent Alterations in Meiotic Recombination Cause Chromosome Segregation Errors in Spermatocytes. *Cell* **171**, 601-614.e13 (2017).

3. D. Jain, *et al.*, rahu is a mutant allele of Dnmt3c, encoding a DNA methyltransferase homolog required for meiosis and transposon repression in the mouse male germline. *PLOS Genet.* **13**, e1006964 (2017).

4. C.-Y. Lee, *et al.*, Mechanism and Regulation of Rapid Telomere Prophase Movements in Mouse Meiotic Chromosomes. *Cell Rep.* **11**, 551–563 (2015).

5. T. Premkumar, *et al.*, Genetic dissection of crossover mutants defines discrete intermediates in mouse meiosis. *Mol. Cell* **83**, 2941-2958.e7 (2023).

6. J. Abramson, *et al.*, Accurate structure prediction of biomolecular interactions with AlphaFold 3. *Nature* **630**, 493–500 (2024).

7. W. Liu, *et al.*, IBS: an illustrator for the presentation and visualization of biological sequences. *Bioinformatics* **31**, 3359–3361 (2015).
